# A fungal effector targets the chloroplast to support biotrophy by balancing disease and plant health

**DOI:** 10.64898/2026.03.23.713812

**Authors:** Djihane Damoo, Matthias Kretschmer, Karen Thulasi Devendrakumar, Sherry Sun, Madeline Iseminger, Leon Pierdzig, Volker Lipka, Kerstin Schmitt, Oliver Valerius, Jennifer Geddes-McAlister, Gerhard H. Braus, Xin Li, Kai Heimel, James W. Kronstad

**Author notes:** Co-corresponding authors: M. Kretschmer,; J. Kronstad. These authors contributed equally and should be considered co-first authors (Djihane Damoo, Matthias Kretschmer).

## Abstract

Fungal pathogens are responsible for substantial crop losses worldwide. There is a pressing need to develop crops with improved disease resistance, especially given that climate change and human activities are exacerbating crop diseases. Our understanding of the molecular mechanisms by which fungi cause disease is incomplete. To address this limitation, we employed proteomics to identify candidate effector proteins from the pathogenic fungus *Ustilago maydis* that co-purified with the chloroplasts of maize host plants during infection. We specifically characterized the role of one putative chloroplast-associated effector, UmPce3, using heterologous expression in the non-host plant *Arabidopsis thaliana*. We discovered that UmPce3 interacts with the chloroplast DEAD-box RNA helicase, AtRH3. Phenotypes associated with the expression of UmPce3 in *Arabidopsis* mirrored those of plants with impaired AtRH3 function and included interference with chloroplast assembly, an impact on photosynthesis, and altered resistance to biotic and abiotic stresses. Support for RH3 as a bona fide effector target was obtained by identifying parallel phenotypic influences of UmPce3 in maize and by demonstrating an interaction between UmPce3 and maize ZmRH3b, an ortholog of AtRh3. Notably, UmPce3 contributes to biotrophy by promoting the virulence of *U. maydis* on maize seedlings and dampening virulence in plants challenged with salinity as an abiotic stress. Overall, this work highlights the chloroplast as a target of fungal pathogenesis and identifies RH3 as a potential hub for pathogen manipulation of organelle function to balance fungal proliferation and host health in support of biotrophy.

**Short summary:** The chloroplast plays a key role in plant immunity, in addition to its central contributions to photosynthesis, metabolism, and tolerance of abiotic stresses. The effector UmPce3 of the maize pathogen *Ustilago maydis* targets the DEAD-box RNA helicase RH3 in host plants to manipulate chloroplast function and enhance fungal pathogenesis. Unexpectedly, UmPce3 also influences host tolerance to salt stress thereby balancing the plant response to biotic and abiotic stressors in support of biotrophic development.

## Introduction

Plants are vulnerable to a wide range of pathogens including viruses, bacteria, fungi, oomycetes, and nematodes. The impact of these pathogens on crop production is estimated to be approximately $220 billion (US) annually, a value that is expected to increase with climate change (Graziosi et al., 2019; Singh et al., 2023). To mitigate threats to food security, it is essential to understand plant immunity so that robust crops with durable resistance can be developed. It is also critical to achieve a deeper understanding of the molecular mechanisms employed by plant pathogens to cause disease.

Upon the initial invasion of host plants, conserved pathogen-associated molecular patterns (PAMPs) are recognized by the plant’s pattern recognition receptors (PRRs) to trigger PAMP-triggered immunity (PTI) (Jones and Dangl, 2006). PTI activation induces mitogen-activated protein kinase (MAPK) cascades, Ca^2+^ fluxes, activation of defense-related genes, reactive oxygen species (ROS) production, and phytohormone signaling (DeFalco and Zipfel, 2021; Su and Gassman, 2023). Pathogens deploy effector proteins to promote disease by a variety of mechanisms, including interfering with plant immune responses, such as PTI (Jones and Dangl, 2006; Khan et al., 2016; DeFalco and Zipfel, 2021). In some cases, effectors can be directly or indirectly recognized by resistance proteins (R proteins) leading to activation of effector triggered immunity (ETI) (Jones and Dangl, 2006; Khan et al., 2016). Therefore, the characterization of effector proteins is a key step to understand the plant processes targeted by pathogens during infection and to enable the use of R proteins for developing resistant crops (Ahmed et al., 2018).

Biotrophic and hemi-biotrophic fungal pathogens deploy effectors to manipulate their hosts with the goal of maintaining plant health sufficient to support pathogen proliferation (Lanver et al., 2017; Schuster et al., 2024; Leiva-Mora et al., 2024; Bhaskar et al., 2025). The fungus *Ustilago maydis* has emerged as a useful model biotrophic pathogen for studying effectors due to the ease of genetic manipulation of the yeast form, and facile virulence testing with maize (*Zea mays)* seedlings (Olicón-Hernández et al., 2019). *U. maydis* is a dimorphic basidiomycete that infects maize or teosinte (*Zea mexicana*) to complete the sexual phase of its life cycle. Interestingly, the fungus incites tumors on any above ground part of the plant and those that form on developing ears are an edible delicacy (Valverde et al., 1995). During the biotrophic stages of its sexual life cycle, *U. maydis* produces 450 to 500 putative effector proteins, most of which lack annotated domains (Ma et al., 2018; Bhaskar et al. 2025). To date, the molecular functions of only few of these effectors have been characterized (Bhaskar et al. 2025). For example, Cmu1 is thought to divert chorismate metabolism into the phenylpropanoid pathway and away from the chloroplast-localized biosynthesis of the plant immunity hormone salicylic acid (SA) (Djamei et al., 2011; Han et al., 2019). Chloroplasts are compelling potential targets of effectors given their central role in plant defense, acting both as environmental integrators and metabolic hubs for the biosynthesis of fatty acids, amino acids, defense phytohormones, and other plant defense-related metabolites (Jelenska et al., 2007; Djamei et al., 2011; Rodriguez-Herva et al., 2012; Kretschmer et al., 2019; Xu et al., 2019; Littlejohn et al., 2020; Breen et al., 2022; Rui et al. 2025; Sun et al., 2025).

We previously found that chloroplast and photosynthetic functions are impaired during *U. maydis* infection (Kretschmer et al., 2017). Additionally, we showed that maize lines impaired in chloroplast development due to a defect in *whirly1* were more susceptible to *U. maydis* infection. These plants have reduced levels of chloroplast ATP synthase, photosystems I and II, cytochrome *b*6f complex and Rubisco (Prikryl et al., 2008; Krupinska et al. 2014; Kretschmer et al., 2017). In the present study, we extended our analysis of *U. maydis* pathogenesis by employing mass spectrometry to identify putative effectors in chloroplasts isolated from *U. maydis*-infected maize seedlings. We prioritized the characterization of one effector designated UmPce3 (*U. maydis* putative chloroplast effector 3) based on an *in silico* prediction of a chloroplast localization signal peptide (cTP) and the finding that a ***Δ****umpce3* deletion mutant had reduced virulence on maize. When expressed heterologously in the non-host *Arabidopsis thaliana*, UmPce3 provoked a curly leaf phenotype, compromised the development of siliques, and increased susceptibility to a biotrophic pathogen of *Arabidopsis*. Subsequent immunoprecipitation and mass spectrometry experiments revealed that UmPce3 interacts with the chloroplast DEAD-Box RNA helicase AtRH3 in *Arabidopsis*. AtRH3 in *Arabidopsis* and its two homologues in maize play crucial roles in ribosome assembly in chloroplasts and in photosynthesis via group II intron splicing of chloroplast transcripts. The *Arabidopsis* lines expressing UmPce3 additionally showed chloroplast dysfunction similar to *Atrh3* knockdown mutants including abnormal chlorophyll content, and altered chloroplast gene expression. Notably, UmPce3 conditioned resistance to abiotic stress imposed by salt thereby maintaining plant health in support of biotrophic development. Parallel phenotypic studies in maize with *U. maydis* mutants lacking UmPce3 supported the conclusion that the effector targets the ZmRH3 homologue to interfere with chloroplast function and promote biotrophic disease.

## Results

### *U. maydis* alters the chloroplast proteome and delivers candidate effectors during infection

Chloroplasts represent prime targets for manipulation of host defense responses and primary metabolic functions during microbial pathogenesis (Littlejohn et al., 2020; Breen et al., 2022; Rui et al. 2025; Sun et al., 2025). In this regard, chloroplast functions are known to be down-regulated during tumor formation induced by *U. maydis* (Kretschmer et al., 2017). Therefore, we performed a proteomics analysis of chloroplasts from infected leaves to identify putative effectors associated with the organelle. We isolated chloroplasts from non-infected control and infected plants at three time points (3, 5, and 7-days post inoculation (dpi)) and analyzed both the chloroplast proteome and the chloroplast-associated proteins from *U. maydis* (Supplemental Table S1). To this end, a 85% PF-Percoll gradient was used to isolate chloroplasts, and enriched chloroplasts from two interfaces of the gradient were combined for protein identification (Fig. 1A). In contrast to chloroplast samples from uninfected plants or infected plants at days 3 and 5 (not shown), the organelles from infected tissue at day 7 showed a green bottom phase in the Percoll gradient thus indicating chlorophyll leakage (Figure 1A, B).

**Figure 1.**
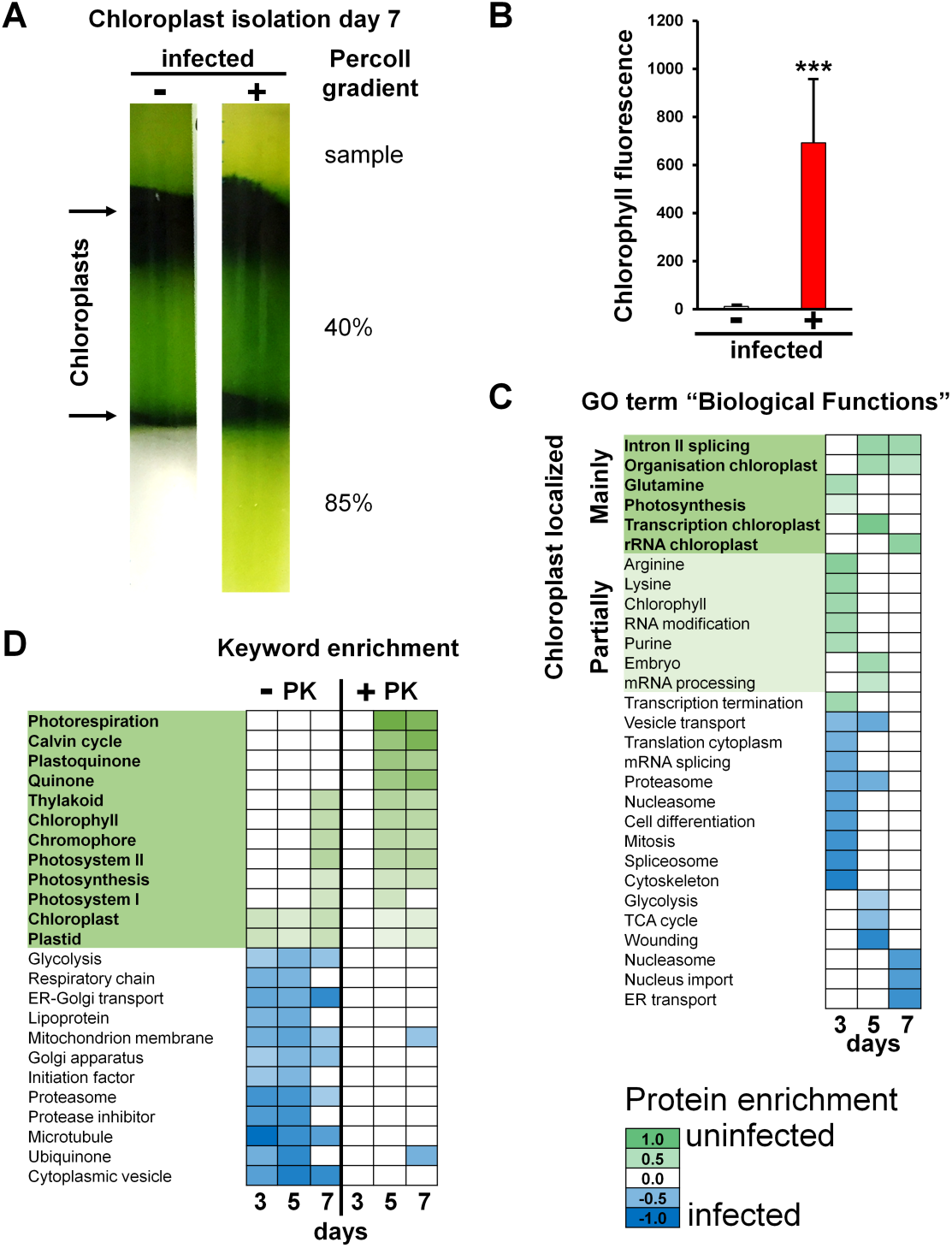
Isolated chloroplasts show altered proteomes during seedling infection with *U. maydis*. A) Chloroplasts from *U. maydis* infected and uninfected maize leaves of the variety Early Golden Bantam were isolated at days 3, 5 and 7, and loaded on a 40% and 85% PF-Percoll step gradient. After centrifugation, the chloroplasts were collected from the 2 interfaces marked with arrows and used for protein isolation. Infected samples at day 7 showed significant chlorophyll leakage in the bottom phase of the step gradient. Chlorophyll leakage was not observed for any uninfected or infected sample at 3 and 5 d post inoculation (dpi). B) Chlorophyll fluorescence was measured for the bottom layers of the gradients for the infected and uninfected samples shown in A, with an excitation of 355 nm and an emission measured at 680 nm. The average with standard deviation was determined. A t-test was used to analyze the data for statistically significant differences with p=***≤0.001. C) GO term enrichments are shown for biological functions of maize chloroplast proteins during the time course of infection at 3, 5 and 7 d for uninfected and infected maize leaf samples. The samples were not treated with proteinase K (PK). Green indicates enrichment in abundance in uninfected samples, while blue indicates enrichment in infected samples. The complete GO term list can be found in Supplemental Table S3. Functions labelled with dark green are exclusively localized to the chloroplast, while functions shaded in light green can partially be found in chloroplasts but are also localized in other cellular compartments. D) The top and bottom 12 enriched keywords are indicated after proteinase K (PK) treatment compared to untreated samples for 3, 5 and 7 dpi of uninfected and infected samples. Green indicates enrichment in abundance in uninfected samples, while blue indicates enrichment in infected samples. The complete keyword list can be found in Supplemental Table S3. Functions labelled in dark green are exclusively localized to the chloroplast. Proteinase K treatment was performed on a second set of samples to reduce non-chloroplast protein and peptide sources and thus to enrich chloroplast-localized protein abundance.

This result suggests compromised structural integrity of the chloroplasts during infection at the later timepoint, a finding consistent with unsuccessful attempts to isolate chloroplasts at 10 dpi.

A second set of chloroplast samples (from 3, 5 and 7 dpi) was treated with proteinase K to remove surface-associated cytoplasmic plant and fungal proteins and to increase the relative abundance of chloroplast-localized proteins. Mass spectrometry analysis of the isolated proteins was performed to identify both the maize and the *U. maydis* proteins for proteinase K treated and untreated samples compared to samples from uninfected leaves. After filtering, a total of 3737 and 395 unique proteins corresponding to maize and *U. maydis*, respectively, were identified in chloroplasts in at least one sample group (Supplemental Tables S1, S2). A total of 88 of the identified *U. maydis* proteins were predicted to be secreted (Supplemental Table S2). Because proteinase K treatment might further compromise chloroplast integrity, the analysis of the chloroplast proteome was initially carried out with enzyme-untreated samples. For the maize proteins, a Gene Ontology (GO) term analysis for biological functions was performed with significant over and under-represented proteins in the chloroplast samples during infection. Enriched proteins in uninfected samples during the time course represented six GO terms unique to chloroplasts as well as seven additional GO terms associated with chloroplasts but also found in other cellular compartments (Supplemental Table S3). These functions were also enriched in the infected samples either at one or all tested time points. Functions related to chloroplast transcription and RNA processing were especially over-represented in the uninfected samples. Furthermore, an enrichment analysis for keywords confirmed that chloroplast functions were enriched after proteinase K treatment in uninfected samples, while non-chloroplast-functions were depleted in the infected samples (Figure 1C, D; Supplemental Table S3).

As mentioned, the proteome analysis also identified *U. maydis* proteins in the proteinase K-treated and -untreated samples (Supplemental Table S2). All of the identified fungal proteins from treated or untreated samples were examined to identify candidate chloroplast-associated effectors. In particular, the fungal proteins identified from proteinase K-treated samples were predicted to be enriched for chloroplast-localized proteins. A total of 125 proteins (105 unique) were identified as differentially abundant among the day 3, 5 and 7 samples, of which 48 (32 unique) were also present in the published modules of candidate *U. maydis* effectors (Lanver et al. 2018). An *in-silico* prediction of chloroplast localization was performed for all identified *U. maydis* secreted proteins using LOCALIZER (https://localizer.csiro.au/) to identify chloroplast Transit Peptide (cTP) motifs (Sperschneider et al., 2017). Five candidate effector proteins were enriched in the chloroplast samples and showed a cTP; each of these were previously identified as effectors including the Stp1 protein required for disease (Table 1, Supplemental Table 2 (full matrix analysis)) (Lanver et al. 2018; Ludwig et al, 2021). Three additional effectors contained a predicted cTP, but were not differentially abundant in any of the conditions. Next, we calculated the delta of the protein intensity of infected minus uninfected samples at the different time points for proteinase K and untreated samples and the average was determined for each of the conditions. The delta protein abundance values were compared with the effector gene expression during infection according to Lanver et al. (2018) to correlate protein data with *in-vivo* gene expression (Supplemental Figure S1). Interestingly, the previously characterized effector Cmu1 (UMAG_05731; Djamei et al. 2011) that modifies chloroplast metabolism was also enriched in protein samples from infected plants. Its abundance increased after proteinase K treatment, suggesting that it may be localized within chloroplasts. However, because Cmu1 lacks a canonical chloroplast targeting signal, additional studies are required to determine whether it is truly localized inside chloroplasts or simply resistant to proteinase K degradation (Supplemental Table S2).

**Table 1:** List of candidate effectors with predicted chloroplast transit peptides (cTPs)

| Gene ID | Function | Secreted protein | Chloroplast localized | LOCALIZER score | cTP position | Detection (day) |
| --- | --- | --- | --- | --- | --- | --- |
| *UMAG_05928*<br>(*UmPce3*) | Candidate effector | Yes | Yes | 1.000 | 87-118 | 3(p)<sup>a</sup> |
| *UMAG_05803* | Acid phosphatase | Yes | Yes | 0.998 | 26-58 | 3(p), 5, 7 |
| *UMAG_02475*<sup>b</sup> | Stp1 | Yes | Yes | 0.987 | 23-67 | 5, 7, 5(p) |
| *UMAG_03860* | Peptidase A1 | Yes | Yes | 0.972 | 32-53 | 3(p) |
| *UMAG_05995* | Undefined | Yes | Yes | 0.637 | 24-84 | 3(p) |

<sup>a</sup>(p)=proteinase K treated samples. <sup>b</sup>Ludwig et al., 2021

As seen in Figure 2A-C and Supplemental Table S2, the protein encoded by UMAG_05928 showed the most significant enrichment of the five effectors in the proteinase K treated samples, especially at days 3 and 5. This result was consistent with the highest gene expression for UMAG_05928 seen at day 4 during infection (Lanver et al. 2018). The abundance of transcripts for UMAG_05928 was confirmed by qPCR with infected samples compared to mating cells grown in culture medium (Figure 2C). We designated the UMAG_05928 product as the candidate effector UmPce3 and selected it for further analysis based on its enrichment in isolated chloroplasts as well as the high probability of a cTP.

**Figure 2.**
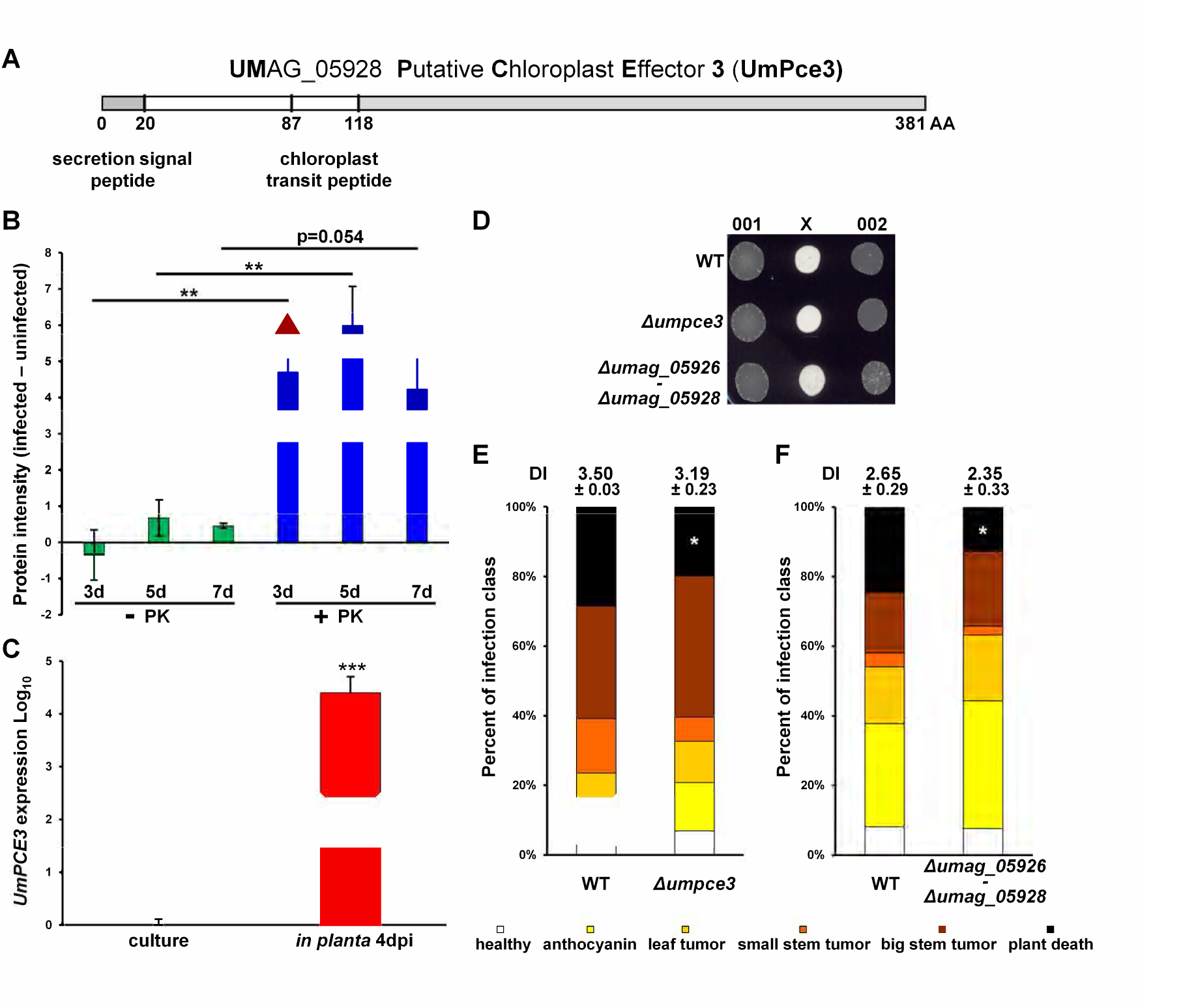
The chloroplast-associated candidate effector UmPce3 is important for fungal virulence of *U. maydis*. A) Diagram of the polypeptide of the candidate chloroplast effector UmPce3. The first 20 amino acids represent the signal peptide for secretion. The predicted chloroplast transit peptide (cTP) is located between amino acids 87 and 118. B) The average protein intensity in infected minus uninfected chloroplast samples for 3, 5 and 7 dpi treated with proteinase K (PK) or untreated. The standard deviation is shown and the data were analyzed with ANOVA plus Tukey’s (p≤0.01 **). The red triangle depicts the timepoint and treatment where the protein was detected as differently abundant during proteomics. The list of all detected *U. maydis* proteins is in Supplemental Table S1 and 2. C) Transcript levels analyzed with qPCR for *UmPCE3* during maize infection at 4 dpi compared to axenic mating culture. The average of 3 biological repeats plus standard deviation is shown. The data were analyzed with a t-test (*** p ≤ 0.001). D) Mating of mutants lacking UmPce3 and the putative effector cluster containing genes UMAG_05926-UMAG_05928 after 48 h at RT in the dark on DCM plus charcoal. Mating between the WT strains 001 and 002 is shown as a control. E) The impact of UmPce3 on virulence of *U. maydis* on maize seedlings of the variety Early Golden Bantam was assessed. The plants were grown under optimal conditions and 7 d old seedlings were infected with mating cultures in the 001 and 002 strain backgrounds with or without deletion of *Umpce3*. Symptoms were scored two weeks after inoculation. The average disease index of 3 biological repeats with standard deviation is shown. The data, including the individual symptom categories, were analyzed with a t-test for statistical significance (* p ≤ 0.05). F) The impact of the candidate effector cluster *ΔUMAG_05926-UMAG_05928* on virulence of *U. maydis* was tested on maize seedlings of the variety Early Golden Bantam. The plants were grown under optimal conditions and 7 d old seedlings were infected with mating cultures in the 001 and 002 strain backgrounds with or without deletion of *ΔUMAG_05926-UMAG_05928*. Symptoms were scored two weeks after inoculation. The average disease index of 3 biological repeats with standard deviation is shown. The data including the individual symptoms categories were analyzed with a t-test for statistical significance (* p ≤ 0.05).

### Loss of UmPce3 reduces the virulence of *U. maydis*

The UmPce3 candidate effector is conserved among smut fungi, and the sequence after the predicted cTP showed the highest conservation in several regions of uncharacterized function (Figure 2A, Supplemental Figure S2A). Initially, the UMAG_05928 gene encoding UmPce3 was deleted in the mating compatible strains 001 (*a2 b2*) and 002 (*a1 b1*) to analyze its contribution to *U. maydis* virulence (Figure 2D, E; Supplemental Figure S3). The UmPce3 deletion strains showed no mating defects that would preclude pathogenicity (Figure 2D). Under optimal plant growth conditions, deletion of the *Umpce3* gene resulted in a modest reduction in the virulence of *U. maydis* in maize seedlings (Figure 2E). Notably, the category of seedling death was significantly reduced upon infection with the deletion strains (black bars in Figure 2E).

The gene encoding UmPce3 was found in a cluster of three genes that included the paralogs UMAG_05926 (e-value: 3e^−16^) and UMAG_05927 (e-value: 8e^−20^). An alignment of the polypeptide sequences is shown in Supplemental Figure S2B. To assess potential redundancy, the full cluster was deleted in the compatible mating strains and the mutants were tested for virulence (Figure 2F). The deletion of the cluster resulted in a similar reduction in virulence as seen for the deletion strain lacking UmPce3, thus indicating that the other paralogs did not contribute to virulence. Consistent with this conclusion, the transcripts for UMAG_05926 and UMAG_05927 were less abundant during infection compared to UMAG_05928 with 23.9-fold lower and 77.3-fold lower expression at day 4, respectively (Lanver et al., 2018). Additionally, the proteins encoded by UMAG 05926 and UMAG 05927 were not detected in the chloroplast proteome analysis. Finally, UMAG_05927 does not contain a predicted chloroplast localization transit peptide, while UMAG_05926 has both predicted chloroplast and a mitochondrial transit peptide as well as a nuclear import signal. Overall, the mutant analysis revealed that only UmPce3 made a significant contribution to the virulence of *U. maydis* in maize seedlings.

### Heterologous expression of UmPce3 in *Arabidopsis* alters plant morphology and impairs plant immunity

We next investigated the function and interaction targets of UmPce3 by expressing a codon-optimized version of the protein in the non-host model plant *Arabidopsis thaliana* and in *Nicotiana benthamiana* (Saado et al., 2022). *Arabidopsis* was employed to make use of available molecular genetic tools relative to the more limited tools for maize. The first macroscopic phenotype observed in the transgenic lines of *Arabidopsis* (Columbia ecotype) was a curled and rolled leaf morphology (Figure 3A). Additionally, the transgenic lines displayed a delay in flowering time, a reduced number of mature seed pods (flower sterility) as well as lower numbers of seeds per mature pod (Supplemental Figure S4A-C). These results indicated an impact of UmPce3 on plant morphology and development when heterologously expressed in *Arabidopsis*. As presented below, the phenotypic impact of UmPce3 was subsequently investigated in the *U. maydis-*maize interaction prompted by phenotypes observed in transgenic *Arabidopsis* lines expressing UmPce3.

**Figure 3.**
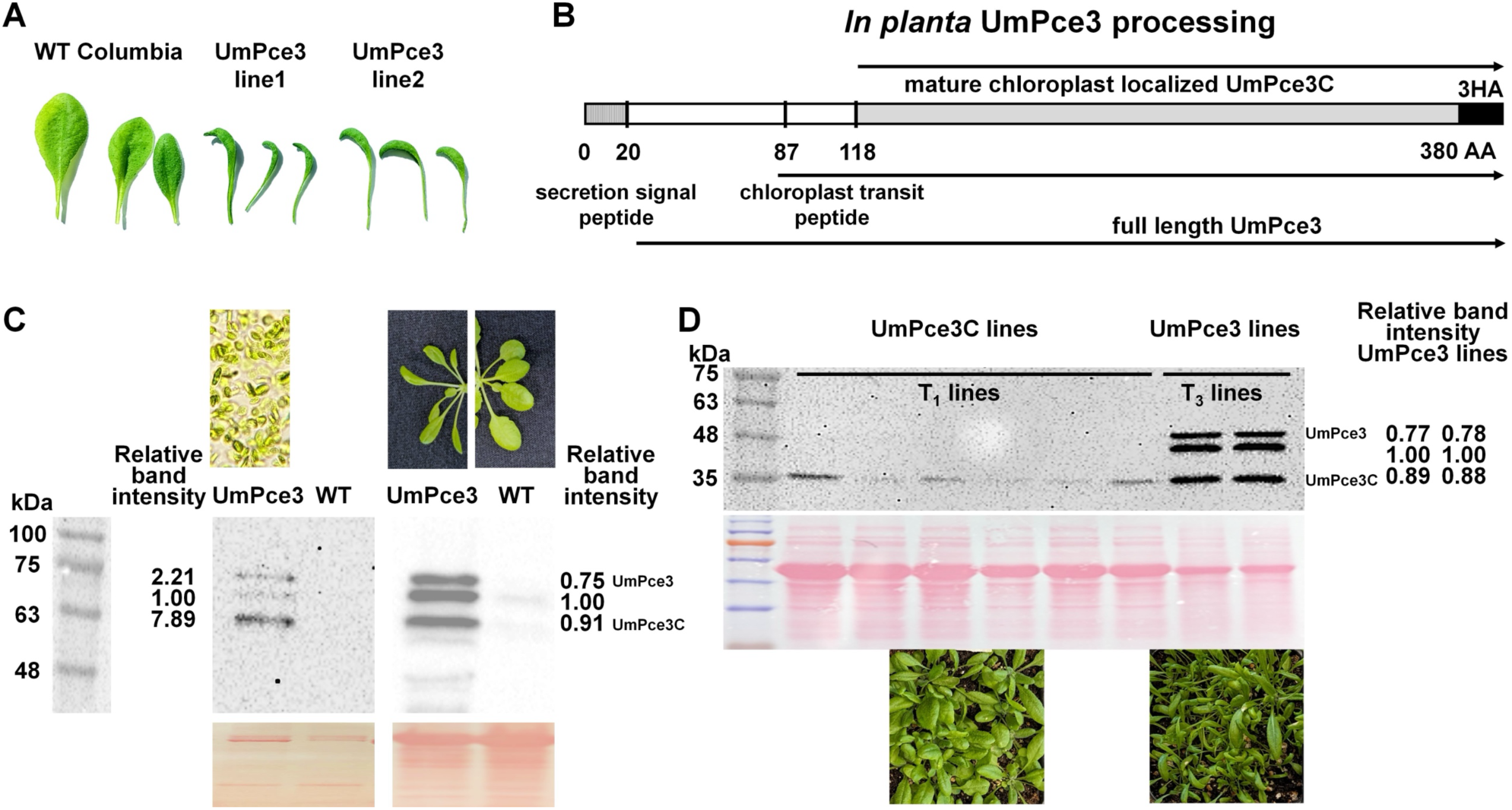
The effector UmPce3 is proteolytically processed and the cTP is important for the morphological impact on *Arabidopsis*. A) Curled and rolled leaf morphology of two independent UmPce3 expressing lines (Line 1: Pce3-2HA-mGFP5 and Line 2: Pce3-mNEON-HA) compared with the WT Col-0 leaves. Lines 1 and 2 were used for all subsequent experiments. B) The proposed proteolytic processing of UmPce3 (UMAG_05928) is depicted. The full-length polypeptide without the fungal secretion signal peptide is designated UmPce3, while UmPce3C corresponds to the mature, chloroplast-localized candidate effector polypeptide. C) Western blot analysis of whole plants and isolated chloroplast samples of *Arabidopsis* plants expressing UmPce3 (2HA-mGFP5). A WT plant is shown for comparison with the altered morphology of the transgenic plant. Chloroplasts were isolated the same way as reported for the maize chloroplasts (Figure 1, Materials and Methods). Pictures of the isolated chloroplasts and the whole plants are shown for reference. The chloroplasts were not proteinase K treated and thus contain minor extra-chloroplast protein contaminants. A ponceau stain was used as loading control (bottom). The protein band intensity ratios of the 3 bands for the plant expressing UmPce3 and for the isolated chloroplasts are shown beside the blots. For the ratios, the center band was used as a reference set to 1. D) Western blot analysis of whole plant protein extracts of *Arabidopsis* plants expressing UmPce3-3HA or UmPce3C-3HA. The smallest protein band observed in UmPce3 corresponds to the protein band observed in plants expressing UmPce3C (lacking the cTP) from the construct pBASTA-35S-UmPce3noctp-3HA. The first six lanes are independent protein samples from whole T1 *Arabidopsis* plants expressing UmPce3C, while the last 2 lanes are T3 plants expressing UmPce3. The protein band intensity ratio of the 3 bands for the UmPce3 expressing lines is shown beside the blot and the plant morphologies are shown below the ponceau stain loading control. For the ratios, the center band was used as a reference set to 1. The same proteolytical processing was independently observed for the HA, mNeonGreen and GFP tagged UmPce3 proteins.

We next tested a role for UmPce3 in suppression of plant immunity given that some effectors in *U. maydis* have this activity and loss of UmPce3 in *U. maydis* led to a reduction in virulence on maize seedlings (Lanver et al., 2018; Bhaskar et al., 2025) (Figure 2E, F). The status of immunity for *Arabidopsis* is routinely assessed with the biotrophic oomycete pathogen *Hyaloperonospora arabidopsidis* (*Hpa*) Noco2 (Thulasi Devendrakumar et al., 2019). Upon challenge with *Hpa* Noco2, two independent lines expressing UmPce3 displayed significantly higher numbers of conidiospores compared to the Columbia (Col-0) ecotype control plants, and the susceptibility of the transgenic plants was similar to that of the positive control *eds1* mutant (Supplemental Figure S4D). However, no differences in the levels of the chloroplast-related plant defense hormone salicylic acid (SA) were observed between the lines expressing UmPce3 and the Col-0 plants (Supplemental Figure S4E). Taken together, we conclude that UmPce3 impacts plant development and leaf morphology, and impairs plant immunity against a biotrophic pathogen but does not compromise SA biosynthesis.

### UmPce3 is proteolytically processed in *Arabidopsis thaliana*

Proteins destined for import into chloroplasts are processed and only the mature functional protein is present in the organelle. Therefore, processing of UmPce3 would be anticipated upon heterologous expression of the protein and import into chloroplasts. To test this idea, two independent *Arabidopsis* lines expressing UmPce3 translationally fused to 3X hemagglutinin tag (3HA) were examined for the size of the protein by immunoblot analysis. Three protein bands were observed with the smallest presumably corresponding to the mature protein localized in chloroplasts (Figure 3B, C). *Arabidopsis* plants expressing UmPce3 consistently showed the three bands with a ratio of ∼0.8: 1.0: ∼0.9, with the middle band used as a reference set as 1. The same proteolytic pattern was independently observed for different UmPce3 proteins tagged at the C terminus with HA and GFP or mNeonGreen (Figure 3C, image on the right of proteins from whole plants; data not shown for UmPce3-mNeonGreen), and the plants expressing these proteins showed the curled and rolled leaf morphology. When isolated chloroplasts were used for protein extraction instead of whole plants, the observed ratio of the three protein bands established for UmPce3 changed to 2.2: 1: 7.9 (left images of isolated chloroplasts and immunoblot, Figure 3C). As the samples were not proteinase K treated and thus still contained some extra-chloroplast protein contaminants, all three protein bands from the whole plant samples were found, but an enrichment of the mature UmPce3C in the isolated chloroplasts was observed (Figure 3C). A modified construct for the expression of a protein corresponding to the mature polypeptide without the chloroplast import signal (designated UmPce3C) was used to confirm the identity of the three protein bands. A single protein band corresponding to the smallest polypeptide was detected in the lines expressing UmPce3C-3HA (Figure 3D). Interestingly, none of the >100 transgenic plants expressing UmPce3C showed the curly leaf phenotype when the chloroplast import signal was missing, thus suggesting a link between the phenotype and chloroplast localization. In combination, these results indicated proteolytic processing of UmPce3 in *Arabidopsis* consistent with import into chloroplasts and that the function of the mature protein is dependent on the chloroplast import signal.

### UmPce3 interacts with the chloroplast-localized DEAD-box-ATP-dependent RNA helicase RH3 of *A. thaliana* and *Z. mays*

We next used *Arabidopsis* plants expressing epitope-tagged UmPce3 in an immunoprecipitation-mass spectrometry (IP-MS) experiment to identify interacting partner proteins (Figure 4A, Supplemental Table S4). First, IP samples from *Arabidopsis* plants expressing UmPce3-2HA-mGFP5 were compared with samples from plants expressing mGFP5, in triplicates. This approach identified 382 proteins with three LFQ intensity values in at least one of the experiment groups. The set included peptides for UmPce3 that were located primarily in the region of the mature polypeptide after the cTP (Figure 4B, Supplemental Table S4; Supplemental Figures S5, S6). This finding is consistent with the processing of UmPce3 described above. The initial IP-MS experiment was performed with a GFP-tagged version of UmPce3, and a second IP-MS experiment was performed with an HA-tagged version to gain confidence in the identification of true interactors (Figure 4A). The comparative analysis of both data sets reduced the number of interacting proteins to 88 (including UmPce3), of which 47 are known to be chloroplast localized (Figure 4B). The complete dataset is presented in Supplemental Table S4 and includes the data for the detected peptides, the initial proteomics analysis, the data for the specific detailed corresponding proteins including potential chloroplast localization, and the associated GO term enrichment for the proteins. The data for the 47 chloroplast proteins included frequent detection of peptides from the S50 and S30 chloroplast ribosomal proteins, as well as from other chloroplast proteins. The GO term enrichment analysis of the 87 identified plant proteins for biological processes (GOBP) identified photosynthesis (e.g., PORB/A and LHCA4), protein localization to the chloroplast (e.g., HSP70-6/7and CLPC1), plastid organization, translation, gene expression and Group II intron splicing as enriched candidate interacting proteins (Supplemental Table S4). The most enriched protein was AtRH3 (Uniprot # Q8L7S8, At5g26742), a chloroplast localized DEAD-box ATP-dependent RNA helicase important for S30 and S50-dependent chloroplast biogenesis and for intron splicing (especially of chloroplast group-II-introns) (Asakura et al., 2012). In *Arabidopsis* and maize, RH3 coimmunoprecipitated with a subset of chloroplast ribosomal proteins similar to those we observed for interaction with UmPce3 (e.g., S30 and S50 chloroplast ribosomal proteins) (Asakura et al., 2012).

**Figure 4.**
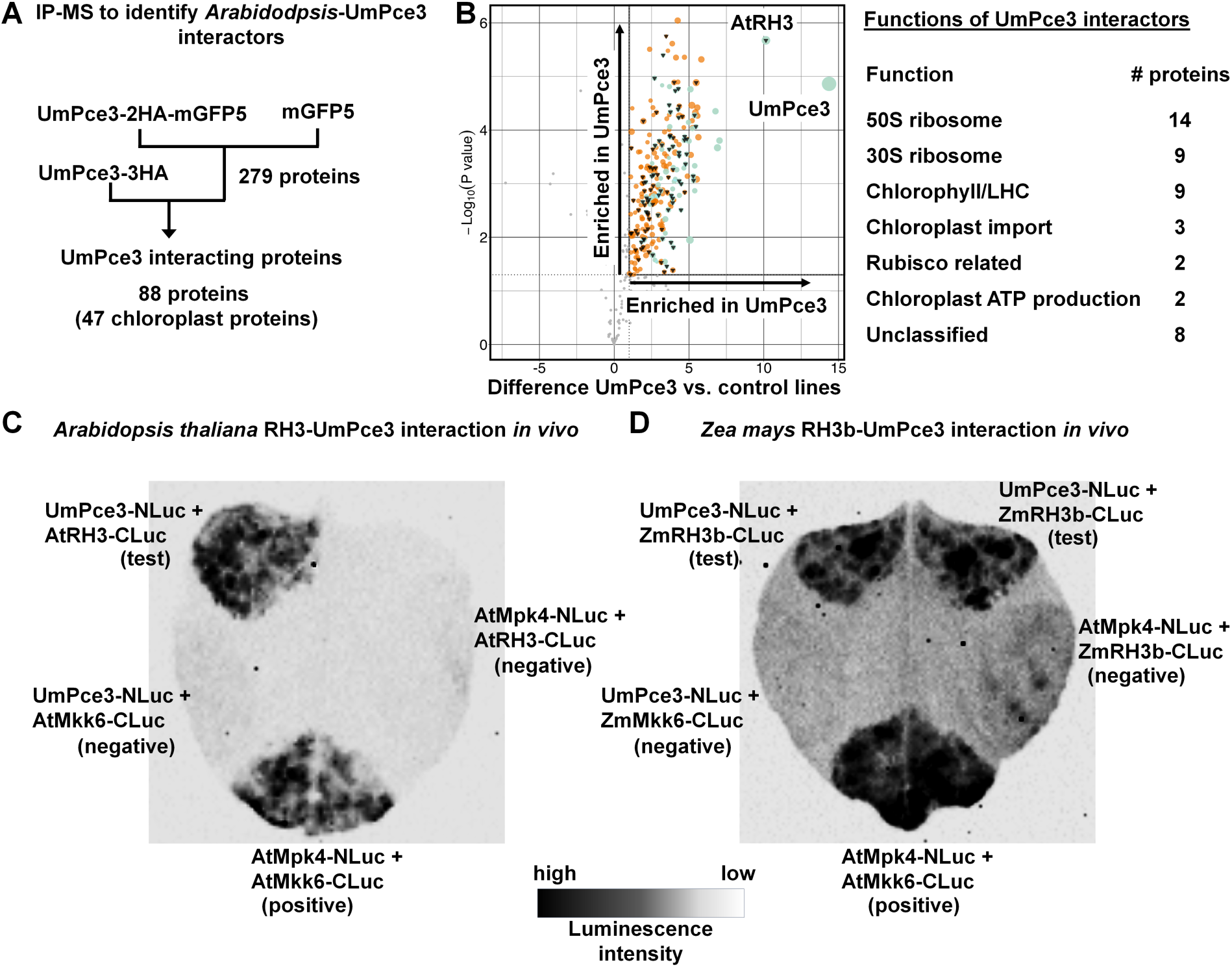
The *U. maydis* effector UmPce3 interacts *in vivo* with the RH3 DEAD-Box RNA helicase from *Arabidopsis* and maize. A) Diagram of the procedure for the identification of UmPce3 interactors. *Arabidopsis* plants expressing GFP-tagged UmPce3 (UmPce3-2HA-mGFP5) or GFP (mGFP5) were used to perform anti-GFP (ChromoTek GFP-Trap® Agarose beads) immunoprecipitation-mass spectrometry (IP-MS) experiments in triplicate. This process identified a total of 382 significant putative interacting proteins with 279 proteins fulfilling the p-value ≤ 0.05 and S_0_ = 1 condition. A second experiment with plants expressing UmPce3-3HA was performed with Pierce™ Anti-HA beads to reduce the number of candidate interactors to 88 proteins including UmPce3, of which 47 were known chloroplast proteins. For more detailed information see Materials and Methods. B) The volcano plot shows the 382 identified putative UmPce3 interactors. A total of 279 significant positive interactors on p value of less than 0.05 were further analysed and marked in the quadrant “enriched in UmPce3” when the Log2 transformed LFQ protein intensity values of the UmPce3 plants compared to GFP expressing plants were p<0.05 according to a t-test, had a value of greater than 1 for t-test differences between those two lines and were detected in a second UmPce3-3HA pulldown experiment. This identified 88 putative proteins including UmPce3. For more detailed information see Materials and Methods. Orange circles show proteins which were only detected in the UmPce3-GFP analysis while turquoise circles show proteins identified in the UmPce3-GFP as well as in the UmPce3-HA analysis. The circle sizes reflect the average MS/MS count for each depicted protein, and chloroplast proteins are marked with a black triangle. The effector UmPce3 (bait) and the putative interactor AtRH3 as best hits are labelled in the volcano plot. A more detailed version of this part can be found in the Supplemental Figure S5. The major functional groups for the 47 chloroplast proteins identified as UmPce3 interactors are listed on the right. C) Split-luciferase assays to confirm the *in vivo* interaction of UmPce3 and AtRH3. Constructs for UmPce3-NLuc and AtRh3-CLuc were transiently expressed by *Agrobacterium tumefaciens* infection of *Nicotiana benthamiana*. 1 mM luciferin was infiltrated and luminescence was recorded according to the depicted scale. A positive control included the known interaction of AtMpk4 and AtMkk6. Negative controls included tested interactions of UmPce3 with AtMkk6, or AtRH3 with AtMpk4. UmPce3 was codon optimized for expression in *N. benthamiana*. D) Split-luciferase assay to confirm the *in vivo* interaction of UmPce3 and ZmRH3b. UmPce3-NLuc and ZmRh3b-CLuc were transiently expressed by *A. tumefaciens* infection of *N. benthamiana*. 1 mM luciferin was infiltrated and luminescence was recorded according to the depicted scale. The positive and negative controls were included as in (C). UmPce3 was codon optimized for expression in *N. benthamiana*.

Given the identification of AtRH3 in *Arabidopsis* as a potential key interactor for UmPce3, we were curious whether ZmRH3 in maize also interacted with UmPce3. ZmRH3 in maize is encoded by two closely related genes, GRMZM2G415491/Zm00001eb211400 (ZmRH3a) and AC198418.3_FG005/Zm00001eb063300 (ZmRhH3b). We employed a split-luciferase assay to test for *in vivo* interactions of UmPce3 with the *Arabidopsis* AtRH3 protein and one of the maize RH3 proteins (ZmRH3b) (Figure 4C, D). *Agrobacterium tumefaciens* was used for transient expression of codon optimized AtRH3-CLuc or ZmRh3b-CLuc with UmPce3-NLuc in *N. benthamiana* leaves. Based on luminescence intensity, *in vivo* protein interactions were observed for the positive control (AtMpk4-NLuc and AtMkk6-CLuc) as well for interactions of AtRH3 and ZmRH3b with UmPce3, while the negative controls did not show luminescence (Figure 4C, D). Taken together, these results support the conclusion that the chloroplast protein RH3 is a target for UmPce3 interaction in both *Arabidopsis* and maize.

### Heterologous expression of UmPce3 in *A. thaliana* provokes phenotypes similar to RH3 mutants in *Arabidopsis* and maize

The interaction between UmPce3 and AtRH3 suggests that transgenic plants expressing UmPce3 may exhibit phenotypes similar to those reported for lines with compromised AtRH3 expression, particularly if UmPce3 exerts an inhibitory influence (Asakura et al., 2012). In *Arabidopsis* and maize, deletion of *RH3* is seedling lethal, while reduced expression leads to pale leaves, delayed growth, and defects in flowering and seed setting (Asakura et al., 2012; Gu et al., 2014). As mentioned above, we observed morphological changes in *Arabidopsis* plants expressing UmPce3 similar to those reported for an *Arabidopsis* mutant defective in RH3 including altered leaf morphology, delayed flowering and compromised seed setting (Figure 3). Furthermore, the IP-MS analyses indicated that UmPce3 not only interacted with AtRH3 but also with 30S and 50S ribosomal proteins similar to what has been reported for AtRH3 (Figure 4; Supplemental Table S4)(Asakura et al., 2012). In *Arabidopsis*, AtRH3 is predominantly found in the interface of the 30S and 50S ribosomal complex during chloroplast biogenesis. Reduced expression of At*RH3* in *Arabidopsis* also results in lower protein levels for photosynthesis-related functions of the photosystems I and II as well as the cytochrome b6f complex (Asakura et al., 2012). Similarly, we found that expression of UmPce3 in *Arabidopsis* led to reduced expression of genes encoding photosynthesis-related functions including PSI (PsaA), PSII (PsbA), Rubisco (RbcS and RbcL), TrnK and, to some degree, the gene for AtRH3 (Figure 5A). The same set of genes showed strongly reduced expression in maize tumors induced by *U. maydis* at day 10 of infection (Figure 5B). These genes included Zm*RH3a* and Zm*RH3b* that were 3.57-fold and 5.10-fold down regulated in tumors after 10 d post infection with *U. maydis*, respectively (Figure 5B, C; Supplemental Table S1) (Kretschmer et al., 2017). Additionally, RH3a, RH3b and TrnK also showed reduced protein levels in isolated chloroplasts during *U. maydis* colonisation of maize (especially at 5 dpi; Figure 5C). Interestingly, the observed reduction in transcript levels of the chloroplast genes due to expression of UmPce3 in *Arabidopsis* was less severe compared to the reductions in the transcripts seen in *U. maydis*-induced maize tumors. This indicates that UmPce3 may not be solely responsible for the expression changes and that other additional components influence chloroplast gene expression in tumors in maize. Additionally, it is interesting that the *trnK* transfer(t) RNA is a known target for group II intron splicing by AtRH3. We note that the phenotypic impact may be less severe in lines expressing UmPce3 compared to a line impaired in AtRH3 function, as effectors often show modulating rather than completely inhibitory activities.

**Figure 5.**
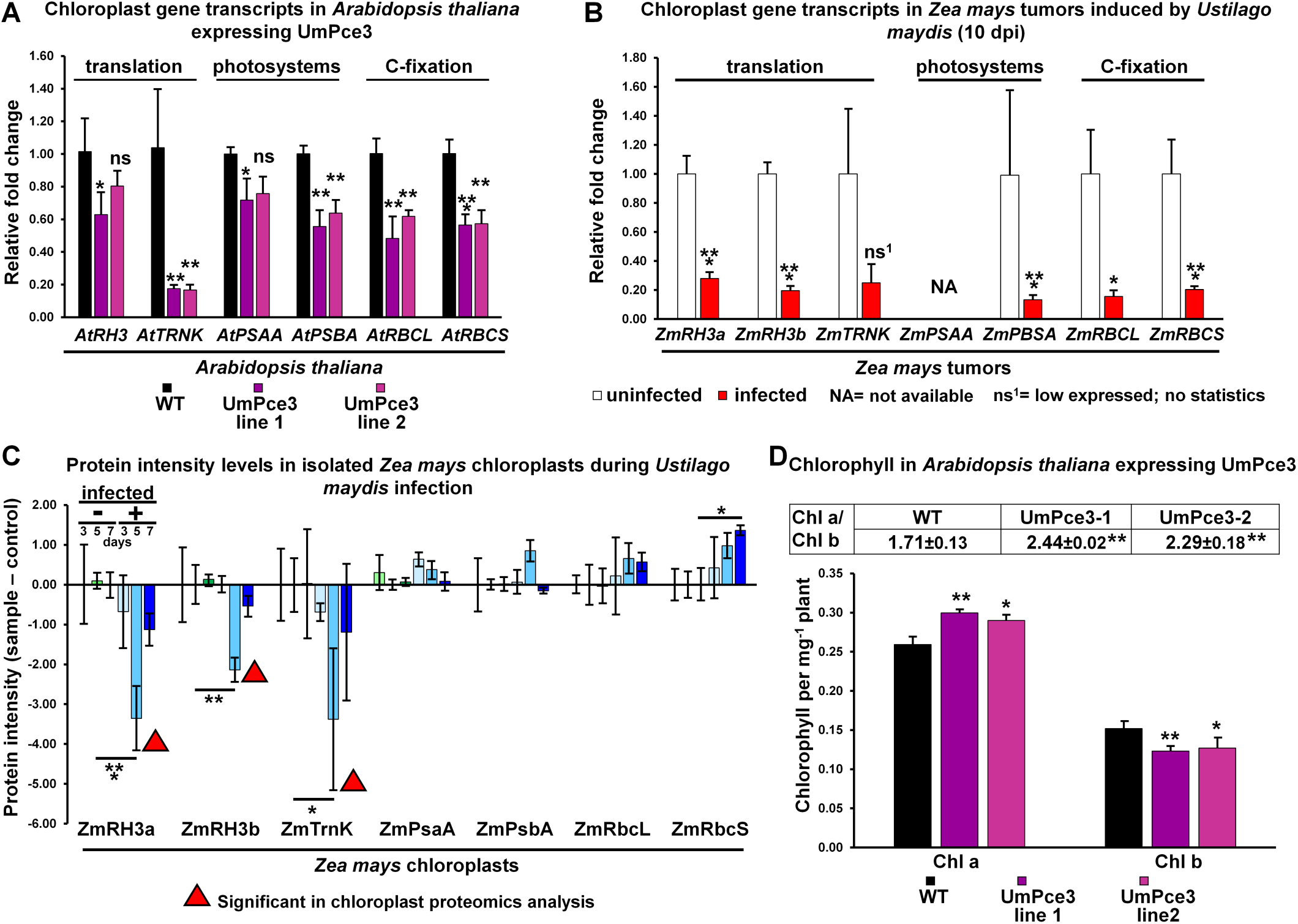
UmPce3 influences chloroplast gene expression in maize tumors and isolated organelles, as well as gene expression and chlorophyll content in *Arabidopsis*. A) Expression of the chloroplast genes in *Arabidopsis* expressing UmPce3 in lines UmPce3-1, UmPce3-2, and in Col-0 WT determined by qPCR. The standard deviations are shown and the data were analyzed with ANOVA plus Tukey’s test (* p=0.05; ** p=0.01; ***p=0.001). B) Expression of chloroplast genes (corresponding genes from A) in maize tumors induced by *U. maydis* 10 dpi. The averages with standard deviation are shown (analyzed as per Kretschmer et al. 2018). The statistical evaluations are according to the RNAseq analysis of Kretschmer et al., (2017b) (* p=0.05; ** p=0.01; ***p=0.001). C) The average protein intensities from the chloroplast proteomics analysis of infected samples minus the uninfected samples at 3, 5 and 7 dpi for the proteins named in A and B. Standard deviations are shown and the data were analyzed with ANOVA plus Tukey’s test (* p=0.05; ** p=0.01; ***p=0.001). The red triangle depicts the timepoint and treatment where the protein was detected as differently abundant by proteomics analysis (Supplemental Table S1). D) Chlorophyll a and b content of three-week old *Arabidopsis* plants expressing UmPce3 in lines Pce3-1, Pce3-2, and in WT Col-0 plants. The average with standard deviation is shown and the data were analyzed with ANOVA plus Tukey’s test (* p=0.05; ** p=0.01).

Besides the changes in chloroplast gene expression, no obvious change in leaf color was observed for *Arabidopsis* lines expressing UmPce3. However, the chlorophyll content in UmPce3-expressing lines was altered with an increase in chlorophyll a and a decrease of chlorophyll b levels (Figure 5D). This led to an overall increase of the chlorophyll a/chlorophyll b ratio compared to Col-0. The observed impact of UmPce3 on chloroplast gene expression, especially of components of the PSI and PSII, led us to test the importance of photosystems for *U. maydis* virulence. Inhibition of PSII by the herbicide Sencor and PSI by Gramoxone both led to increased virulence of *U. maydis* at sub-herbicidal concentrations (Figure 6). Furthermore, PSII inhibition with Sencor led to increased symptoms during infection with no plant toxicity at the tested herbicide dilutions and no direct toxicity for *U. maydis* cells in culture (Figure 6A, B). Inhibition of PSI with Gramoxone resulted in toxicity to *U. maydis* in culture and only led to an impact on fungal virulence close to concentrations with herbicidal activity (Figure 6C, D). Together, these results suggest that both photosystems contribute to the immunity of maize challenged with a biotrophic fungal pathogen, but that PSII appears to have a greater impact.

**Figure 6.**
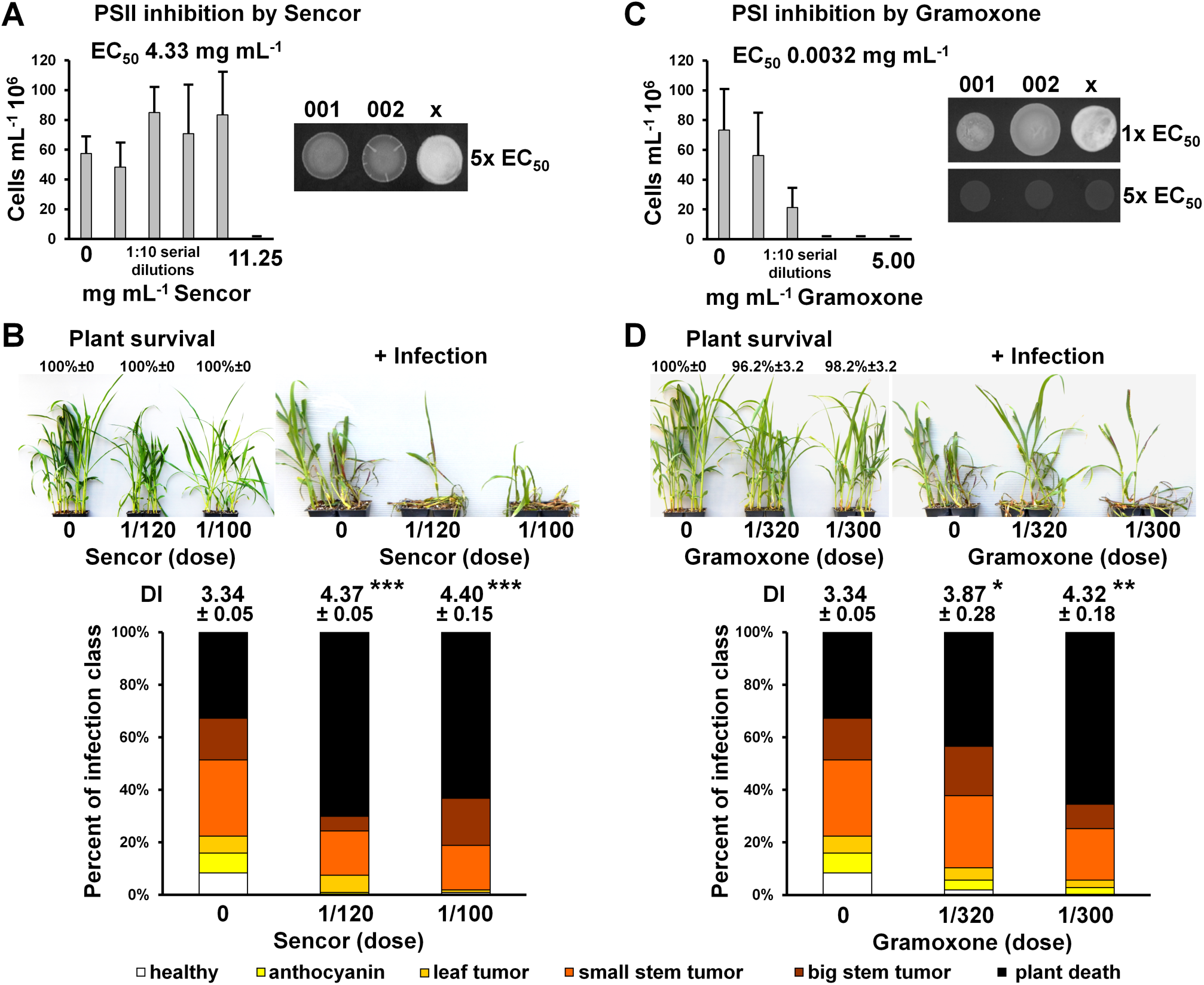
PSII and PSI inhibition compromises maize immunity during colonisation by *U. maydis*. A) Impact of the PSII inhibitor Sencor on *U. maydis* growth in PDB cultures and mating on DCM with charcoal for WT strains 002 (growth curve and mating) and 001 (mating). The WT strain 002 was grown in 5 mL of PDB for 24h at 200 rpm 30°C with an initial inoculum of 10^6^ cells. Sencor was added in 10-fold serial dilutions. The EC50 was calculated, and the EC50 concentration as well as 5-times the EC50 concentration were tested for an impact on mating for 48 h at room temperature. The average with standard deviation is shown. B) Seven-day old seedlings of Early Golden Bantam were sprayed with dilutions of Sencor in water until run off and the seedlings were inoculated 2 d later with a mating mixture of the *U. maydis* WT strains 001 and 002 with a final concentration of 5×10^6^ cell per mL. The symptoms were scored after 2 weeks and pictures of uninfected control plants and infected plants at the different herbicide dilutions were taken. Plant survival was also calculated. The average with standard deviation is shown and the data were analyzed with ANOVA plus Tukey’s test (*** p=0.001). Note that the full recommended dose of Sencor causes the death of all maize seedlings. C) Impact of the PSI inhibitor Gramoxone on *U. maydis* growth in PDB cultures and mating on DCM with charcoal for WT strains 002 (growth curve and mating) and 001 (mating). The WT 002 strain was grown in 5ml of PDB for 24 h at 200 rpm 30°C with an initial inoculum of 10^6^ cells. Gramoxone was added in 10-fold serial dilutions. The EC50 was calculated, and the EC50 concentration as well as 5-times the EC50 concentration were tested for an impact on mating for 48 h at room temperature. The average with standard deviation is shown. D) Seven-day old seedlings of Early Golden Bantam were sprayed with dilutions of Gramoxone in water until run off and the seedlings were inoculated 2 d later with a mating mixture of the *U. maydis* WT strains 001 and 002 with a final concentration of 5×10^6^ cell per mL. The symptoms were scored after 2 weeks and pictures of uninfected control plants and infected plants at the different herbicide dilutions were taken. Plant survival was also calculated. The average with standard deviation is shown and the data were analyzed with ANOVA plus Tukey’s test (*** p=0.001). Note that the 1/20^th^ of the recommended dose of Gramoxone causes the death of all maize seedlings. Additionally, the same untreated plants were used for the Sencor and Gramoxone experiments and are shown in the “0 dose” treatment images in B and D.

### Loss of UmPce3 enhances *U. maydis* virulence on maize experiencing salt stress, and expression in *Arabidopsis* phenocopies the response to abiotic stress

*Arabidopsis* plants compromised for AtRH3 expression have altered metabolism of the plant hormone abscisic acid (ABA) and increased salt sensitivity during seed germination and plant development (Lee et al., 2012). In this context, *Arabidopsis* seeds expressing UmPce3 showed reduced germination rates on medium containing salt compared to Col-0 (Figure 7A). In contrast, two-week-old plants grown on soil showed an inverse salt sensitivity where Col-0 was more sensitive than the lines expressing UmPce3 (Figure 7B). Interestingly, the virulence of *U. maydis* was reduced for WT inoculations while the virulence of the deletion strains lacking UmPce3 was higher in maize seedlings treated with increasing salt concentrations (Figure 7C). In the latter case, a higher proportion of dead seedlings was observed compared to WT or non-salt treated plants. These results suggest that UmPce3, as a chloroplast targeting protein, has a dampening activity in plants experiencing salt stress that may promote the establishment and maintenance of a biotrophic interaction. This activity would be consistent with the dependence of biotrophic pathogens on a living host to propagate and fulfill their lifecycle. Plant death or impaired vitality due to abiotic stress would be contrary to this dependence and UmPce3 may target chloroplast functions to specifically modulate the biotrophic interaction depending on environmental stress conditions.

**Figure 7.**
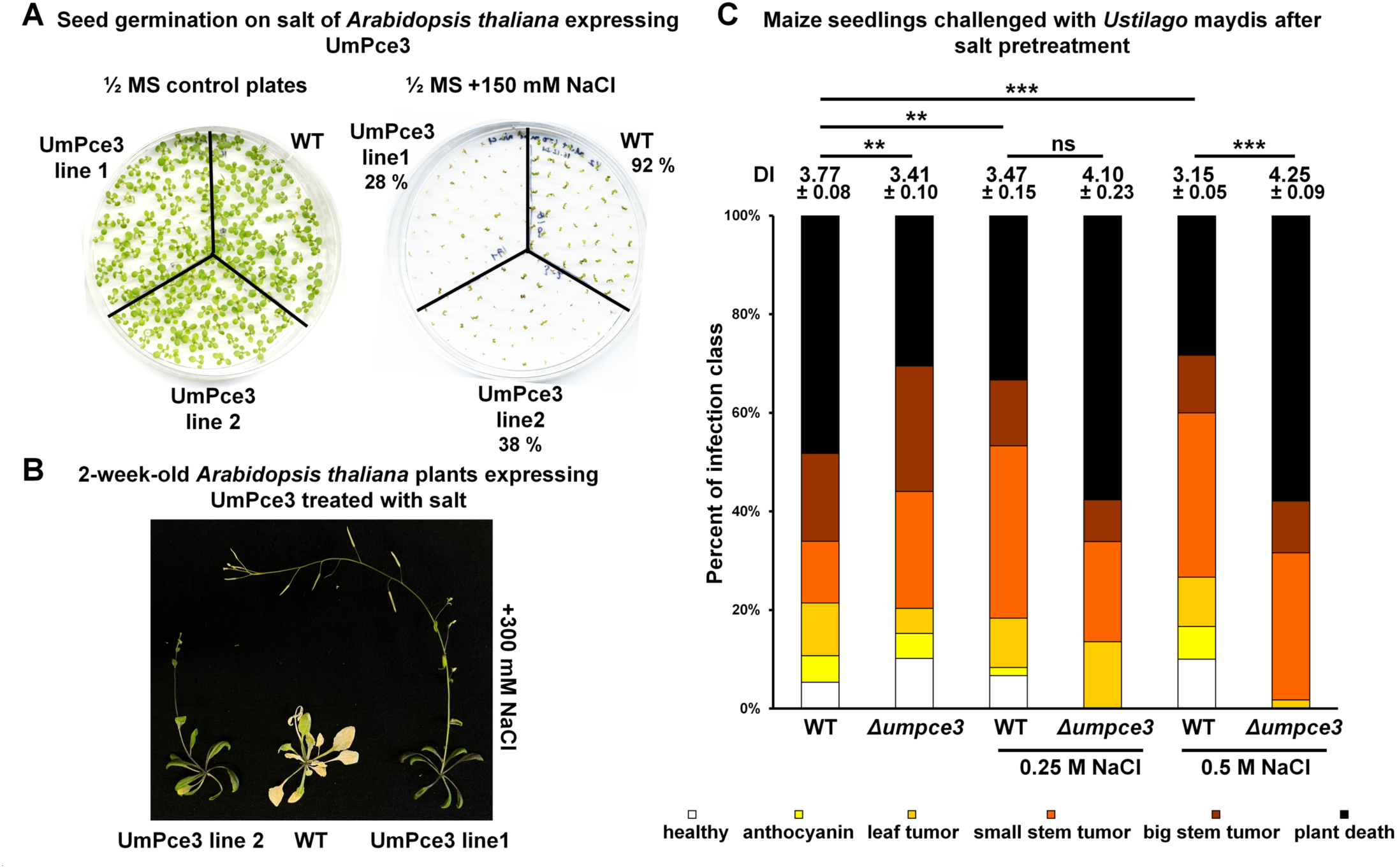
UmPce3 affects the response of *Arabidopsis* and maize to salt stress. A) Percent of germinated seeds of *Arabidopsis* lines expressing UmPce3 on 1/2 MS plates containing 150 mM NaCl. UmPce3 expression lines had a lower germination rate compared to the WT Col-0. Germination on a ½ MS plate without salt is shown as control. B) Arabidopsis lines expressing UmPce3 and comparable WT plants were grown in soil and watered with a solution of 300 mM NaCl. The plants were photographed after 2 weeks. C) Seedlings of the maize variety Early Golden Bantam were grown in sealed planting trays and watered on day 5 post planting with 0.25 M or 0.5 M NaCl solutions, or with water. On day 7 post planting, the seedlings were infected with a mating culture of 001 and 002 with 5×10^6^ cells per ml of the WT mating strains or a strain lacking UmPce3. The plants were watered with water without additional salt during the infection period. After 2 weeks post inoculation, the disease symptoms were scored and the disease index as averages plus standard deviation was calculated. The data were analyzed with ANOVA plus Tukey’s test (ns not significant; ** p=0.01; *** p=0.001).

### UmPce3 impacts retrograde signaling consistent with an interaction with RH3

Retrograde signaling from the chloroplast to the nucleus can be activated when chloroplast function is perturbed, such as upon treatment with the antibiotic lincomycin (Brunkard and Burch-Smith, 2018; Loudya et al., 2024). Interestingly, AtRH3 is also implicated in retrograde signaling in *Arabidopsis* (Asakura et al., 2012). We therefore tested whether UmPce3 has an impact on AtRH3-related retrograde signaling by germinating *Arabidopsis* seeds on media containing lincomycin (Figure 8A). *Arabidopsis* lines expressing UmPce3 showed improved germination and seedling greening compared to Col-0. We also found that the growth and mating of *U. maydis* was not inhibited by lincomycin (Figure 8B). However, maize seedlings challenged with lincomycin and simultaneously infected with WT strains of *U. maydis* showed increased disease symptom formation (Figure 8B). Lincomycin was not lethal for maize at the concentrations tested, but zones of chlorosis and leaf tip browning were observed at higher concentrations. When maize plants were challenged with lincomycin and infected with *U. maydis* mutants lacking UmPce3, an increase in disease symptoms was observed compared to the infections with the WT strains (including a high percentage of dead seedlings). These results again support the idea that UmPce3 exerts a biotrophy-modulating activity in adverse environmental conditions for maize growth, as may result in impaired retrograde signaling.

**Figure 8.**
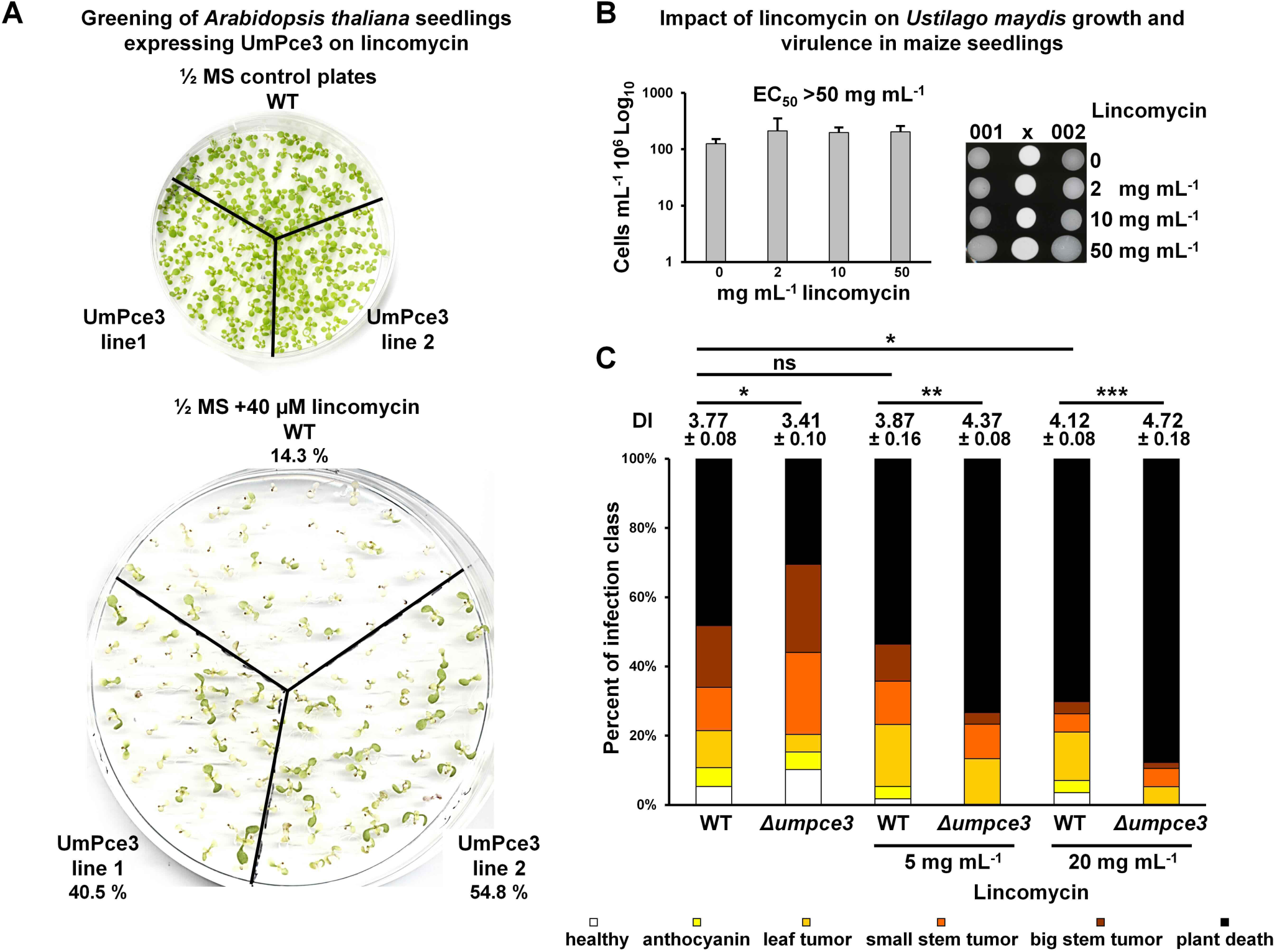
UmPce3 affects retrograde signaling induced by lincomycin in *Arabidopsis* and maize. A) Percent of green seedlings of *Arabidopsis* lines expressing UmPce3 on 1/2 MS plates containing 40 μM lincomycin. Lines expressing UmPce3 had a higher germination rate and seedling greening compared to Col-0. Germination and seedling greening on a ½ MS plate without lincomycin is shown as a control. The control plate was also included with the assays with NaCl treatment (Figure 8A). B) Impact of the antibiotic lincomycin on *U. maydis* growth in PDB cultures and mating on DCM with charcoal for WT strains 002 (growth curve and mating) and 001 (mating). The WT 002 strain was grown in 5 ml of PDB for 24 h at 200 rpm 30°C with an initial inoculum of 10^6^ cells. The EC50 exceeded the maximal tested concentration of lincomycin. The impact of lincomycin on mating was tested on DCM charcoal plates with the same concentrations tested for growth. Plates were grown 48 h at room temperature. The average with standard deviation is shown. C) Early Golden Bantam maize seedlings were infected with *U. maydis* 7 days post planting with WT and UmPce3 deletion strains with and without lincomycin. The seedlings were infected with a mating culture of 001 and 002 with 5*10^6^ cells per mL containing lincomycin for the WT and UmPce3 deletion strains. The lincomycin concentrations were 5 and 20 mg per mL or water only. The disease symptoms were scored at 2 weeks post inoculation, and the disease index as an average plus standard deviation was calculated. The data were analyzed with ANOVA plus Tukey’s test (ns not significant; * p=0.05; ** p=0.01; *** p=0.001). Lincomycin was not lethal for maize, but streaks of chlorosis and leaf tip browning were observed for the assay with 20 mg per mL lincomycin.

### Impaired chloroplast biogenesis interferes with *U. maydis* virulence

We next performed virulence assays with seedlings lacking Zm*RH3b*, one of the two *RH3* paralogs in maize, to investigate the importance of chloroplast biogenesis for the establishment and maintenance of biotrophy and tumor development. The Zm-*rh3b-1* line shows partially impaired chloroplast biogenesis. As a control, we also tested *U. maydis* virulence on a maize line lacking Zm-*ppr53* that has a more severe defect in chloroplast biogenesis (Belcher et al., 2015; Zoschke et al., 2016). Nine-day-old seedlings grown from Zm-*rh3b-1* seeds from segregating ears were genotyped and infected with *U. maydis*. Infection symptoms were scored after two weeks and correlated with the genotype. Large stem tumors at the base of the stems were seen for the majority of the WT/WT or WT/*rh3b-1* plants, but not on *rh3b-1/rh3b-1* plants (Supplemental Figure S7A, B). Small stem tumors that indicated successful biotrophic colonisation were the most severe symptom found in the *rh3b-1/rh3b-1* plants. Infection of these seedlings resulted in a high proportion of dead plants as well as the appearance of a new symptom class characterized by the collapse and death of leaves 3 and 4. The third and fourth leaves are generally the leaves with the most prominent disease symptoms in the seedling assays upon *U. maydis* infection because leaves 1 and 2 have already emerged when the seedlings are inoculated. The majority of Zm-*rh3b-1* seedlings were unable to establish a biotrophic interaction with *U. maydis* as seen in the average disease index of 4.8 vs 4.1 seen for WT. A score of 4 for an infected plant corresponds to large stem tumors at the base of the stem, while 5 represents death of the 3^rd^ and 4^th^ leaves without tumor development. This phenotype was also seen in the Zm-*ppr53* line with a more severe chloroplast defect, where the establishment of a biotrophic interaction leading to tumor development was not observed in any of the seedlings (Supplemental Figure S7). Overall, we conclude that impaired chloroplast biogenesis interfered with the ability of *U. maydis* to establish biotrophic disease. This finding is consistent with an interaction of UmPce3 with ZmRH3 to influence the establishment and maintenance of biotrophy.

## Discussion

Beyond their functions in primary metabolism in plants, chloroplasts also contribute to plant immunity and are therefore prime targets for pathogen effectors. The key defense functions include generating ROS, mediating Ca^2+^ oscillations, and synthesizing the hormones jasmonic acid and SA (Zabala et al., 2015; Sowden et al., 2018; Littlejohn *et al*., 2020; Irieda and Takano, 2021; Kachroo et al 2021; Breen et al., 2022; Rui et al. 2025). These contributions are disrupted by effectors from many types of plant pathogens (i.e., viruses, bacteria, fungi and oomycetes) to promote virulence, as revealed in many recent reports (Jelenska et al., 2007; Rodríguez-Herva et al., 2012; Halane et al., 2018; Kretschmer et al., 2019; Medina-Puche et al., 2020, Gao et al., 2020; Tang et al., 2020; Wang et al., 2021, 2023; Yang et al., 2021; Cao et al., 2024; Fu et al., 2024; Jaswal et al., 2024; Jin et al., 2024; Shang et al., 2024; Song et al., 2024; Xiao et al., 2024; Xu et al., 2024; Zhang et al., 2024; Asghar et al., 2025; Liu et al., 2025). In this study, we employed a proteomics approach to discover that UmPce3 is an effector that targets the chloroplast to influence the biotrophic development of *U. maydis* on maize. UmPce3 therefore joins a growing list of effectors from biotrophs (e.g., rust and powdery mildew fungi) that target chloroplast functions (Djamei et al., 2011; Kretschmer et al., 2019; Wang et al. 2023). For example, effectors from *Puccinia striiformis* f. sp. *tritici* target TalSP, a component of the cytochrome b6/f complex in wheat. This complex functions in photosynthesis and is also targeted by the CSEP80 effector from *Erysiphe nectar* during grapevine infection (Mu et al. 2023). Photosynthetic functions may be a common target in the chloroplast as demonstrated by previous studies with bacterial pathogens (Zabala et al., 2015). For example, HopNI from *Pseudomonas syringae* is a cysteine protease that cleaves PsbQ, a component of photosystem II (Rodriguez-Herva et al., 2012). Given that *U. maydis* influences photosynthesis and provokes chlorosis in developing tumors on maize, we focused on a detailed characterization of UmPce3 to understand the contribution of the protein to disease (Kretschmer et al., 2017).

We found that UmPce3 interacts with the chloroplast protein AtRH3 in *Arabidopsis* along with other proteins suggestive of a specific hub comprised of AtRH3 and proteins for ribosome biogenesis (e.g., 30S and 50S ribosomal subunits). RH3 is a chloroplast DEAD-box RNA helicase characterized in *Arabidopsis* and maize (Asakura et al., 2012). Null mutants in *Arabidopsis* are embryo lethal and characterization of a mutant with a T-DNA insertion in intron 9 (the weak allele At*rh3-4* with reduced *RH3* transcript levels) revealed defects in the splicing of specific group II introns and ribosome biogenesis (Asakura et al., 2012; Lee et al. 2013; Gu et al., 2014). The phenotypes of the At*rh3-4* allele in *Arabidopsis* indicated that RH3 influences chloroplast development and photosynthesis including reduced chlorophyll content resulting in pale green seedlings. In particular, the mutant has reduced accumulation of subunits for PSI, PSII and the cytochrome b6/f complex. Consistent with an interaction with AtRH3, the lines expressing UmPce3 displayed phenotypes reminiscent of an AtRH3 knockdown including the altered chlorophyll content and salt sensitivity (Asakura et al., 2012). When expressed in *Arabidopsis*, UmPce3 provoked a number of phenotypes including an abnormal leaf morphology, altered chlorophyll content, salt sensitivity, reduced PSI and PSII and increased susceptibility to *Hpa* Noco2. We also observed reduced transcript levels of the *trnK* tRNA for the amino acid lysine in the UmPce3 expressing lines.

The information on the role of AtRH3 in *Arabidopsis* and the identified phenotypes of leaf morphology, immune dysfunction and biotic stress sensitivity provoked by UmPce3 guided parallel studies in maize. That is, we hypothesized that UmPce3 interacts with the two maize ZmRH3 proteins affecting the splicing of intron-containing genes in the chloroplast and phenotypes such as abiotic stress sensitivity. Both of the ZmRH3 proteins were detected along the developmental leaf gradient of expanding maize leaves with expression peaking in the sink-source transition. This expression pattern is interesting because the infected leaf undergoes a shift from source to sink during a *U. maydis* infection (Horst et al., 2008). ZmRH3s are maximally expressed in the sink-source transition zone of expanding maize leaves (Asakura et al., 2012) and localize to chloroplasts, stroma, thylakoids and nucleoids. Notably, the *whirly* gene product (ZmWhy1) that regulated chloroplast development is found in the same locations, has a function in chloroplast ribosome biogenesis, and contributes to susceptibility to *U. maydis* (Prikryl et al., 2008; Asakura et al., 2012; Kretschmer et al., 2017a; Ren et al., 2017). It was previously shown that both Zm*RH3* paralogs are downregulated approximately three to five-fold at 10 days post infection (Kretschmer et al., 2017b). We also observed this downregulation in our proteomics data (Supplemental Table S1). ZmRH3 paralogs in maize are known to co-sediment with 50S pre-ribosomal subunits in a sucrose gradient and ZmRH3 also associates with Group II introns and pre-50S ribosomal particles *in vivo* (Asakura *et al.,* 2012). Overall, these results are consistent with the idea that ZmRH3b and ZmWhy1 may be components of a critical chloroplast hub for organelle transcription and translation that is exploited by *U. maydis*. Mutants lacking these proteins are more susceptible to the fungus. Loss of ZmPRR53, a pentatricopeptide repeat protein that binds chloroplast RNAs (Zoschke et al., 2016), also increases susceptibility thus providing further support for the chloroplast as an effector target. A possible connection between RH3 with plant immunity is reinforced by the recent discovery that the protein plays a role in antiviral defense by loading small RNAs onto Argonaute for silencing during viral infections (Huang et al., 2025).

As a proposed hub component, RH3 may also be targeted by pathogens to influence retrograde signaling from the chloroplast to the nucleus (Loudya et al., 2024). Evidence for a link between RH3 and retrograde signaling comes from the observation that At*RH3* transcripts and protein are elevated in mutants lacking the caseinolytic protease (Clp) core complex (Asakura et al., 2012). Mutants with defects in the protease are deficient in chlorophyll and are impaired in chloroplast rRNA processing. Asakura et al. (2012) proposed that the increase in AtRH3 compensates for loss of the protease and is due to signaling from the chloroplast to the nucleus to influence transcription of the nuclear At*RH3* gene. In support of the idea that *U. maydis* infection influences retrograde signaling, we identified an impact of lincomycin on maize susceptibility to the fungus. Lincomycin inhibits protein synthesis in chloroplasts and interferes with retrograde signaling (Brunkard and Burch-Smith, 2018). Furthermore, RNA helicases are involved in different steps of RNA metabolism from ribosome biogenesis to RNA export and RNA decay, and it is therefore possible that UmPce3 interacts with a protein complex that contains ZmRH3 leading to a broader influence on chloroplast and cellular functions (Bourgeois et al., 2016). These findings suggest the need for a more comprehensive functional characterization of RH3 and associated proteins with regard to pathogen attack, chloroplast biology and broader cellular functions such as retrograde signaling.

Overall, expression of UmPce3 in *Arabidopsis* caused similar phenotypes to those of the At*rh3-4* mutant. Consistent with the phenotype of AtRH3 knockdown or T-DNA insertion mutations in *Arabidopsis*, maize seedlings experiencing salt stress had decreased susceptibility to infection by WT *U. maydis*. Interestingly, the salt-stressed plants had increased susceptibility to the *Δumpce3* mutant suggesting an interaction between the activity of the effector and abiotic stress. We hypothesize that this interaction is also mediated via ZmRH3 because of known roles of chloroplasts and DEAD box helicases in the plant response to abiotic stress (Gu et al., 2014, Lee et al., 2012). For example, the difference in salt susceptibility of the germinating seeds versus two-week old plants expressing UmPce3 may be related to the AtRH3-dependent accumulation of abscisic acid (ABA) during salt stress at different stages of plant development. Salt stress is known to induce ABA accumulation in *Arabidopsis* while ABA catabolism is a crucial step to break seed dormancy and trigger germination (Jia et al., 2002; Ali et al., 2019). Other factors mediate the cross talk between chloroplast and abiotic stress (Breen et al., 2022). For example, the SA receptor non-expressor of pathogenesis-related genes 1 (NPR1) protein can localize to chloroplasts in response to salt stress, and lincomycin influences its subcellular localization (Jayakannan et al., 2015; Seo et al., 2020; 2023; Zavaliev and Dong, 2024). Overall, these observations suggest that UmPce3 impacts chloroplast function in a manner that supports biotrophic development by maintaining plant health in the presence of abiotic stress.

The proposed impact of UmPce3 on ZmRH3 and the chloroplast is summarized in Supplemental Figure S8 to emphasize the influence of the effector on the maintenance of plant health in response to abiotic stress and on *U. maydis* virulence to support biotrophic development. To our knowledge, RH3 and retrograde signaling have not yet been identified as targets of known effectors. Our results suggest that RH3 and associated proteins may represent a hub targeted by effectors to influence organelle biogenesis and retrograde signaling between the chloroplast and the nucleus. Therefore, effectors from other pathogens may target this hub. Additionally, other identified candidate effectors from *U. maydis* that target the chloroplast await characterization. We surmise that these other chloroplast effectors or nuclear induced regulation of chloroplast functions contribute to the biotrophy of *U. maydis* because loss of UmPce3 had a modest impact on virulence, although it is possible that redundancy with ZmRH3a buffered the response to the fungus.

## Materials and Methods

### Growth conditions for *Ustilago maydis* and construction of deletion strains

The *U. maydis* strains were grown in potato dextrose broth (PDB) or double complete medium (DCM) with 1% glucose at 30°C and 220 rpm (Holliday, 1974). Mating was tested on DCM plates with 1% activated charcoal at room temperature in the dark. An overlap PCR approach was used to construct deletion mutants in the 001 (518, *a2b2*) and 002 (521, *a1b1*) strain backgrounds. Upstream and downstream flanking regions of the gene for UmPce3 (UMAG_05928) or the gene cluster (UMAG_05926 to UMAG_05928) were amplified from genomic DNA of strain 002 and combined during an overlap PCR reaction with the hygromycin B resistant marker amplified from plasmid pIC19RHL (Wang et al., 1988). The final construct was amplified with nested primers and used for biolistic transformation according to Toffaletti et al. (1993). All primers can be found in Supplemental Table S5. Transformations were carried out on DCM plates with 0.95 M sorbitol. Selection of the deletion strains was performed on DCM plates with 250 μg/mL hygromycin B and genotypes were verified by PCR and genome hybridization. For genomic hybridization to verify *Δumpce3* mutants, 25 ug of RNase-treatedDNA was digested with the restriction enzymes EcoRI and HindIII. The digested DNA was separated on a 0.8% agarose gel and the DNA was transferred with 20X SSC capillary blotting on to a Hybond™- N+ membrane (GE Healthcare, Mississauga, ON, Canada). Hybridization was carried out with the DIG DNA labeling and Detection starter Kit II (Roche Canada, Mississauga, ON, Canada) according to manufacturer’s recommendations and detection was performed with the ChemiDoc™ XRS+ system from Bio-Rad and ImageLab version 4.0.1 software (Bio-Rad, Mississauga, ON, Canada) (Supplemental Figure S3).

### Chloroplast isolation, proteinase K treatment and proteomics analysis

Maize seedlings (∼100 plants per replicate) of the variety Early Golden Bantam were grown for one week and infected at day seven with a mating culture of the WT strains 001 and 002 at a final concentration of 10^7^ cells per mL to achieve robust disease symptoms. Leaf areas with visible infected areas on the 100 inoculated plants were selected at days three, five or seven after infection and corresponding leaf areas of uninfected plants were also harvested. The total weight for each biological repeat was between 7 and 8 g. The infected leaf areas were lightly macerated in a mortar and pestle in 100 mL of ice-cold chloroplast isolation buffer (50 mM HEPES-KOH pH 8, 330 mM sorbitol, 2 mM EDTA pH 8, 5 mM ascorbic acid, 5 mM cysteine, 0.05% BSA). Subsequently, the leaves were further macerated in a Ninja pulse blender (Ninja bl300c) with 10 s bursts (repeated three times). The samples were filtered through two layers of cheese cloth, and the filtrate was centrifuged for three min at 1300xg at 4°C. The samples were then processed for chloroplast isolation and loaded onto a Percoll step gradient as described by van Wijk et al., 2007. Intact chloroplasts enriched at the 40% and 85% interfaces were harvested with a transfer pipette and pooled. The chloroplasts were washed three times (50 mM HEPES-KOH pH 8, 330 mM sorbitol, 2 mM EDTA pH 8), pelleted and flash frozen.

A proteinase K treatment was performed with a second set of chloroplast samples isolated at the same timepoints to eliminate protein contaminations from *U. maydis* and other non-chloroplast sources and to enrich for chloroplast-localized proteins. The chloroplasts were incubated for 20 min in 1 mL isolation buffer with 200 μg per mL proteinase K on ice before loading onto the step gradient. The samples from the step gradients were then processed the same way as the non-treated chloroplast samples. The concentration of proteinase K employed lysed the chloroplasts at ∼40 min on ice. No visible lysis was detected at 20 min for uninfected samples. We noted that the chloroplasts of infected samples from 7 d showed chlorophyll leakage without proteinase K treatment compared to uninfected samples. No obvious chlorophyll leakage was observed for either uninfected or infected samples harvested at earlier times. Thus, chloroplasts appear to be structurally weakened during infection especially at later timepoints and therefore proteinase K treatment might be disadvantageous for overall comparison of the chloroplast proteins during an infection time course. We therefore focused our analysis primarily on the samples not treated with proteinase K and evaluated the treated samples from early timepoints specifically to identify *U. maydis* proteins associated with the chloroplasts and to eliminate other highly abundant extra-chloroplastic fungal proteins.

Total proteins of the isolated chloroplast samples were extracted as previously described (Ball and Geddes-McAlister, 2019). Briefly, chloroplasts were briefly washed with PBS, resuspended in 100 mM Tris-HCl, pH 8.5 with a proteinase inhibitor capsule (PIC) and mechanically disrupted by probe sonication in an ice bath. Next, 2% (final) sodium dodecyl sulphate (SDS) was added, followed by reduction with 10 mM dithiothreitol (DTT) at 95°C for 10 min at 800 rpm, alkylation with 55 mM iodoacetamide (IAA) at room temperature in the dark, and acetone precipitation (80% final) at -20 °C overnight. Precipitated proteins were pelleted, washed twice with 80% acetone, and air-dried prior to solubilization in 8 M Urea and 40 mM HEPES. Endoproteinase Lys-C (LysC) was added at 1:100 µg protein and incubated for 3 h at room temperature, followed by addition of 300 µl 50 mM ammonium bicarbonate (ABC) and 1:100 µg protein of trypsin with overnight incubation at room temperature. Protein concentration was determined by the BSA tryptophan assay (Wis̈niewski and Gaugaz, 2015). Digestion was stopped by addition of 10% v/v trifluoroacetic acid (TFA) per sample and the acidified peptides were loaded onto Stop-And-Go extraction tips (StageTips) (containing three layers of C_18_) for desalting and purification according to the standard protocol (Rappsilber et al., 2007).

### Mass spectrometry measurement

Samples were eluted from StageTips with 80% acetonitrile (ACN)/0.5% acetic acid, dried with a SpeedVac concentrator, and peptides were resuspended in sample buffer (2% ACN and 0.1% TFA). Approximately 2 µg of protein was subjected to nanoflow liquid chromatography on an EASY-nLC system (Thermo Fisher Scientific, Bremen, Germany) on-line coupled to an Q Exactive HF quadrupole orbitrap mass spectrometer (Thermo Fisher Scientific, Bremen, Germany). A 50 cm column with 75 μm inner diameter was used for the chromatography, in-house packed with 3 μm reversed-phase silica beads (ReproSil-Pur C_18_-AQ, Dr. Maisch GmbH, Germany). Peptides were separated and directly electrosprayed into the mass spectrometer using a linear gradient from 5% to 60% ACN in 0.5% acetic acid over 150 min at a constant flow of 300 nl/min. The linear gradient was followed by a washout with up to 95% ACN to clean the column. The QExactive HF was operated in a data-dependent mode, switching automatically between one full-scan and subsequent MS/MS scans of the fifteen most abundant peaks (Top15 method), with full-scans (*m*/*z* 300–1650) acquired in the Orbitrap analyzer with a resolution of 60,000 at 100 *m*/*z*.

### Mass spectrometry data analysis and statistics

Raw mass spectrometry files were analyzed using MaxQuant software (version 1.6.10.43) (Cox and Mann, 2008). The derived peak list was searched with the built-in Andromeda search engine (Cox et al., 2011) against the reference *U. maydis* proteome (downloaded from UniProt, Dec. 2024; 6,805 sequences) and *Zea mays* proteome (downloaded from UniProt Dec. 2024; 63,237 sequences). The parameters were as follows: strict trypsin specificity was required with cleavage at the C-terminal after K or R, allowing for up to two missed cleavages. The minimum required peptide length was set to seven amino acids. Carbamidomethylation of cysteine was set as a fixed modification (57.021464 Da) and N-acetylation of proteins N termini (42.010565 Da) and oxidation of methionine (15.994915 Da) were set as variable modifications. Peptide spectral matching (PSM) and protein identifications were filtered using a target-decoy approach at a false discovery rate (FDR) of 1%. ‘Match between runs’ was enabled with a match time window of 0.7 min and an alignment time window of 20 min. ‘Split by taxonomy ID’ was enabled; relative label-free quantification (LFQ) of proteins was performed using the MaxLFQ algorithm integrated into MaxQuant using a minimum ratio count of 1, enabled FastLFQ option, LFQ minimum number of neighbors at 3, and the LFQ average number of neighbors at 6 (Cox et al., 2014).

Statistical analysis of the MaxQuant-processed data was performed using the Perseus software environment (version 1.6.2.2)(Tyanova et al., 2016). Hits to the reverse database, contaminants, and proteins only identified by site were removed. LFQ intensities were converted to a log scale (log_2_), and only those proteins present in triplicate within at least one sample set were used for further statistical analysis (valid-value filter of 3 in at least one group). Missing values were imputed from a normal distribution (downshift of 1.8 standard deviations and a width of 0.3 standard deviations). A Student’s *t*-test was performed to identify proteins with significant differential abundance (*p-*value ≤ 0.05) (S_0_ = 1) between samples employing a 5% permutation-based FDR filter (Benjamini and Hochberg, 1995).

### Plant materials and growth conditions

*Arabidopsis thaliana* and *Nicotiana benthamiana* plants were grown in Sunshine® Mix #4 soil in a growth chamber with a light dark cycle of 16h to 8h at 22 °C. For constitutive heterologous expression of UmPce3 (UMAG_05298) in *Arabidopsis,* a GENEWIZ-codon-optimized UmPce3 sequence was synthesized and cloned by GENEWIZ into the pBASTA-35S-3HA vector. The plasmids and primers for preparing transformation constructs are listed in Supplemental Table S5. For all *Arabidopsis* expression lines, the gene for a codon optimized version of UmPce3 was transcribed from the constitutive 35S promoter and the open reading frame lacked the secretion signal peptide but contained C-terminal fusions with the HA epitope, GFP, or mNeonGreen. Lines UmPce3-1 and UmPce3-2, with constitutive expression of UmPce3, correspond to lines 17-1 (Pce3-2HA-mGFP5) and 3-2 (Pce3-mNEON-HA), respectively. The pBASTA-35S-UmPce3noctp-3HA construct was used to express the UmPce3 protein lacking the chloroplast targeting sequence (designated UmPce3C). The Pce3C (Pce3noctp-3HA) construct was amplified by PCR using the primers listed in Supplementary Table S5 and cloned downstream of the CaMV 35S promoter via KpnI/SpeI double digestion. The resulting plasmid was sequence-verified and introduced into *Agrobacterium tumefaciens*. The floral dip method described by Clough and Brent (1998) as used to transform *Arabidopsis* with *A. tumefaciens* strain GV3101 carrying the described vectors Supplemental Table S5. Seeds from the transformed plants were selected on half strength Murashige and Skoog medium (1/2 MS) containing 0.6% agar, 0.5% sucrose, and 15 mg/L Basta for selection of transgenic seedlings of the T1 generation. Protein extracts were prepared from multiple independent T1 transgenic plants. The same Basta selection was used to generate T2 and T3 lines and to evaluate the Mendelian segregation of the offspring.

The two independent transgenic lines (UmPce3-1 and UmPce3-2) were used to investigate developmental differences of *Arabidopsis* plants expressing UmPce3. The time from seeding until the appearance of the first open flower was recorded. For seed production of the different strains, 10 well-developed seed pods for WT and the UmPce3 expression lines were harvested when the pods turned yellow. The expression lines produced fewer well-developed seed pods compared to WT and the best available developed seed pods were used for the assay. The pods were vortexed in 1 mL water with two glass beads in 2 mL screw cap tubes until the seeds were released. The average seed number was determined and calculated as seeds per pod for each plant and each line. To determine the number of mature, well-developed, seed pods, the plants were grown until leaves and seed pods turned yellow. Seed pods were counted as well as the number of undeveloped/aborted flowers.

Chloroplasts isolated from *Arabidopsis* plants expressing UmPce3 (Pce3-2HA-mGFP5) were isolated as for the maize chloroplasts, with the exception that whole leaves were used and the amount of plant material was ∼100 mg. The isolated chloroplasts were used for protein extraction and immune blotting.

### Infection assays with Hyaloperonospora arabidopsidis Noco2

The oomycete *Hyaloperonospora arabidopsidis* Noco2 was used to assess the impact of UmPce3 expression on *Arabidopsis* immunity. Two-week-old *Arabidopsis* plants expressing UmPce3 and control plants were spray inoculated with a solution of 50,000 spores per mL. The plants were transferred to a tray and covered with a transparent plastic cover and incubated at 18°C in a growth chamber. Seven days after infection, the leafy plant parts were harvested in test tubes and the spores were released from the plants by vortexing in 1 mL of water. The total number of spores relative to the fresh weight was determined by spore counting with a Neubauer cell counting chamber.

### Virulence assays with maize seedlings

The maize variety Early Golden Bantam was used for all maize infection assays. Seven-day-old seedlings were infected with 50-100 µl of cultures containing a 1:1 mixture in water of the mating compatible strains in the 001 and 002 background. The final cell density equaled 5×10^6^ cells per mL. Infections assays were conducted in triplicate with approximately 100 plants per inoculation. The disease symptoms were scored at two weeks with each plant assigned the most severe presented symptom ranging from a scale of 0 (no symptoms) to 5 (seedling death) (Kretschmer et al., 2018). This protocol was also used to test symptom formation in plants exposed to salt stress, lincomycin, herbicides, or for infection assays with the maize lines *rh3b-1*/+ X *rh3b-1*/+ and *ppr53-2*/+ X *ppr53-3*/+ (obtained from Dr. Alice Barkan as seeds of ears from segregating plants).

Salt treatment was imposed by watering the plants in sealed trays starting at day five after planting with water without or with 0.25 M and 0.5 M NaCl. Two days later, the seedlings were infected and subsequently only watered with water. To test the impact of retrograde signaling on virulence, a sterile solution of lincomycin was added to the inoculum solution at 5 mg/mL or 20 mg/mL. The seedlings were infected at day 7 and scored after 14 days. The herbicides Gramoxone (Syngenta Crop Protection, Calgary, AB, Canada) and Sencor (Bayer Crop Science, Calgary, AB, Canada) were employed to inhibit PSI or PSII during disease development at sub-herbicidal concentrations. Specifically, Gramoxone was used at 1/300^th^ and 1/320^th^ and Sencor was used at 1/100^th^ and 1/120^th^ of the recommended dose. The herbicides were diluted in water and applied to the seedlings at day 7 until run-off. Infection was performed at day 9 to reduce the impact of the herbicides on *U. maydis*. Symptoms were scored 14 days later. For all compounds, the toxicity for *U. maydis* was tested by growth in PDB medium with increasing concentrations of the compounds, and the EC50 was calculated. An impact on the mating of *U. maydis* cells was tested for the EC50 concentration and 5x the EC50 concentration on DCM agar medium containing charcoal.

The importance of the maize chloroplast protein ZmRH3b for the virulence of *U. maydis* was tested on nine-day old seedlings from segregating ears of *rh3b-1/x* crossed with *rh3-1/x* (170 seedlings in total in 3 biological repeats) and was compared with a Zmppr53 (*Zmppr53-2*/+ X *Zmppr53-3*/+) line with a severely compromised chloroplast developmental phenotype. Maize leaves 1 and 2 are already emerged at day 9 and the infection takes place by injecting the mating culture in the center of the seedling stem, 1 cm above soil level, close to the meristem. Infection symptoms are subsequently observed mainly in developing leaves 3 and 4, after the 14-day infection period. As the chloroplast-impairment phenotype of the Zmrh3b-1 seedlings even after 9 days was not clear, all seedlings were genotyped with primers Rh3b_for and Rh3b_rev for WT and RH3b_for with a 1:1 mixture of primers mu1 and mu2 for transposon integration in ZmRH3b (Supplemental Table S5). The infection phenotype was scored after 14 days and correlated with the plant genotype. The infection phenotype scale from above was amended with a new class 5 (leaves 3 and/or 4 were collapsed and dead) and class 6 represents whole plant death.

### Quantification of transcript levels with qPCR

The transcript level of the gene for UmPce3 in maize seedlings 4 days after infection was compared by qPCR with levels from cells in culture. The infected tissue was ground into a fine powder in a mortar and pestle with sea sand and liquid nitrogen. The RNA was isolated with the RNAeasy Mini Kit according to the manufacturer’s instructions (Qiagen, Toronto, ON, Canada). Total RNA (2.5 μg) was treated with DNAse to remove genomic DNA with the Turbo DNase Kit (Thermo Fischer Scientific, Waltham, MA, USA). Subsequently, 500 ng of DNase-treated total RNA was transcribed into cDNA with oligodT primers and the High-Capacity cDNA Reverse Transcription Kit (Thermo Fisher Scientific, Waltham, MA, USA). Each qPCR reaction contained 2 µl of 1:5 diluted cDNA with gene-specific primers for UmPce3 and syf1 (UMAG_03842) as housekeeping gene. The fold change was calculated with the ΔΔ^Ct^ method. Primers for qPCR are listed in Supplemental Table S5. For *Arabidopsis* lines expressing UmPce3 chloroplast genes transcripts, 200 ng of RNA were reverse transcribed with the gene-specific primers listed in Supplemental Table S5 and amplified using a One-step RT-PCR kit (NEB, Whitby, ON, Canada).

### Determination of chlorophyll a and b in *Arabidopsis* plants expressing UmPce3

The leaves of plants were collected and the fresh weight determined. The samples were ground into a fine powder in a mortar and pestle with liquid nitrogen. The macerated samples were dissolved in 80% acetone, vortexed and incubated at 4°C in the dark for four h. The samples were vortexed once every hour. After incubation, the samples were centrifuged at 10,000 rpm for 5 min to remove solids. The absorbance of the supernatant of each sample was measured at 663 nm for chlorophyll a and at 645 nm for chlorophyll b. The chlorophyll concentrations and the ratio were calculated according to Senthilkumar et al. (2021) and normalized against the fresh weight of the sample.

### Extraction of proteins from *Arabidopsis* and immunoblotting

For the verification of the protein levels of *Arabidopsis* plants expressing UmPce3, total proteins from whole plants were extracted in lysis buffer (0.1% SDS, 2% β-mercaptoethanol, 0.1M Tris, pH 8) after grinding of the tissue in liquid nitrogen in a mortar and pestle. The protein samples were mixed with 4x Laemmli buffer and loaded on a 10% SDS polyacrylamide gel. After separation, the proteins were wet transferred (90 V, 2h) to a polyvinylidene difluoride (PVDF) membrane (GE Healthcare, Boston, MA, U.S.A.). After blocking with Tris-buffered saline with Tween 20 (TBST) and 5% skim milk, the membranes were either incubated with monoclonal anti-HA 1:10000 antibodies (Thermo Fisher Scientific, Waltham, MA, USA) or anti-GFP-horseradish peroxidase antibodies (HRP) (anti-GFP HRP) (Santa Cruz Biotechnology, Dallas, TX, USA), and anti-mouse HRP (Bio-Rad, Hercules, CA, USA) at 1:5,000. The ChemiDoc™ XRS+ system from Bio-Rad and ImageLab version 4.0.1 software was used for detection (Bio-Rad, Mississauga, ON, Canada). One initial colorimetric image of the protein ladder was taken followed by a second image to detect chemiluminescence from the bound antibody. The two pictures were overlaid with ImageLab version 4.0.1 software. A ponceau stain was performed as a loading control.

### Preparation of *Arabidopsis* samples for immunoprecipitation-mass spectrometry

For IP-MS experiments, five-week-old *Arabidopsis* plants expressing UmPce3-2HA-mGFP5, UmPce3-3HA or mGFP5 alone (as control) were grown in short day conditions, harvested and the plant material was ground in a mortar and pestle to a fine powder with liquid nitrogen. The plant material was incubated in extraction buffer (250 mM sucrose, 100 mM HEPES-KOH, 5% glycerol, 10 mM EDTA pH 8, 1 mM DTT, 1% Triton X-100 and 2 X protease inhibitor cocktail) for 30 minutes at 4°C. The samples were then centrifuged to remove solids. Equilibrated ChromoTEK GFP-Trap® Agarose beads (Proteintech®, Rosemont, IL, USA) were incubated with the supernatant overnight at 4°C. For HA-tagged samples the supernatant was incubated with Pierce™ Anti-HA magnetic beads (Thermo Fisher Scientific, Waltham, MA, USA). The beads were washed three times with the extraction buffer and twice with the same extraction buffer without protease inhibitor complex and the proteins eluted in 0.2 M glycine with a pH of 2.5. The samples were then precipitated with chloroform and methanol according to Wessel and Flugge 1984.

The isolated protein samples were suspended in 8 M urea and separated on a 12% SDS-polyacrylamide gel. When the dye front reached halfway the gels were stained with Colloidal Coomassie (Kang et al., 2002). Each lane of the three biological repeats was cut out from the gel and the three gel pieces underwent an in-gel trypsin digestion to generate and release peptides of the proteins (Shevchenko et al., 1996). A StageTip purification was then performed as described above with two layers of C_18_ material (Rappsilber et al., 2007). The resulting peptides were resuspended in sample buffer containing 2% ACN and 0.1% formic acid. Reversed-phase liquid chromatography (LC) was performed on 6 μl of each sample for peptide separation using an RSLCnano Ultimate 3000 system (Thermo Fisher Scientific). Peptides were loaded on an Acclaim PepMap 100 precolumn (100 μm by 2 cm, C_18_, 5 μm, 100 Å; Thermo Fisher Scientific) with 0.07% TFA at a flow rate of 20 μL/min for 3 min. Analytical separation of peptides was performed on an Acclaim PepMap RSLC column (75 μm by 50 cm, C_18_, 2 μm, 100 Å; Thermo Fisher Scientific) at a flow rate of 300 nL/min. The solvent composition was gradually changed over 94 min from 96% solvent A (0.1% formic acid) and 4% solvent B (80% acetonitrile, 0.1% formic acid) to 10% solvent B within 2 min, to 30% solvent B within the next 58 min, to 45% solvent B within the following 22 min, and to 90% solvent B within the last 12 min. All solvents and acids were Optima grade for liquid chromatography mass spectrometry (LC-MS; Thermo Fisher Scientific). Eluting peptides were on-line ionized by nano-electrospray ionization (nESI) using the Nanospray Flex Ion source at 1.5 kV (liquid junction) and transferred into a Q Exactive HF mass spectrometer (Thermo Fisher Scientific). Full scans in a mass range of 300 to 1,650 *m/z* were recorded at a resolution of 30,000 followed by data-dependent top 10 HCD fragmentation at a resolution of 15,000 (dynamic exclusion enabled).

### Mass spectrometry data analysis and statistics

For *A. thaliana* and *U. maydis* protein assignments, the MS/MS2 data from the IP-MS samples were searched against the *A. thaliana* and reference proteome (downloaded from UniProt) using the default MaxQuant (version 1.6.10.43) software settings and performing label-free quantification (Cox and Mann, 2008). The *A. thaliana* proteome fasta file additionally contained the amino acid sequence for UmPce3 with the Uniprot identifier A0A0D1BVZ6. The Perseus software was used (version 1.6.0.7; Tyanova et al., 2016) for statistical analysis as similar as described above for the chloroplast proteome analysis. Only proteins present in triplicate within at least one sample set (UmPce3-GPF or control set) were used for statistical analysis. A Student’s t-test was performed to identify proteins with significant differential abundance (p-value ≤ 0.05) (S_0_ = 1) between samples employing a 5% permutation-based FDR filter (Benjamini and Hochberg, 1995). Proteins were only further considered to be a putative UmPce3 interaction partner if they were also identified with a LFQ intensity value in the UmPce3-3HA sample. This reduced the number of candidates from 279 to 102 proteins. Finally, proteins were only kept and candidates if they were (i) not at all identified in the negative GFP control or (ii) the difference in LFQ intensity between negative GFP control and UmPce3-3HA samples was at least 1. This removed an additional 14 proteins from the list, and 88 remained, including UmPce3 as bait.

### Split-Luciferase assays for assessing interactions of RH3 and UmPce3 proteins

The coding sequence of UmPce3, codon-optimized and synthesized for *N. benthamiana* (GENEWIZ), was cloned into a pCAMBIA-based vector containing the N-terminal coding sequence of luciferase (Vector 2C) using a SpeI/KpnI double-restriction digest. In parallel, AtRH3 was amplified using primers containing DraIII restriction sites, as detailed in Supplemental Table 5 and cloned into the pCAMBIA vector carrying the C-terminal coding sequence of luciferase (Vector 2B) and ZmRh3b (codon-optimized and synthesized for *N. benthamiana* by GENEWIZ) was cloned into the Vector 2B by GENEWIZ. All genes were transcribed from the constitutive CaMV 35S promoter. The constructs were transformed into *A. tumefaciens* strain GV3101 with subsequent growth over night in LB with 15 µg/mL gentamicin, 35 µg/mL rifampicin and 50 µg/mL kanamycin. After centrifugation, the cells were washed with 10 mM MgCl_2_ solution and resuspended at an OD_600_ = 0.5 (for each construct) in infiltration buffer (10 mM MgCl_2_ and 150 µM acetosyringone and 10 mM of MES pH 5.6) in the dark. After induction for 3 h, the *Agrobacterium* cells containing UmPce3 and either AtRH3 or ZmRH3b plus the corresponding controls were infiltrated with a syringe without a needle in the abaxial leaf side of four-week-old *N. benthamiana* leaves. Two days after infiltration, 1 mM luciferin was infiltrated with a syringe without a needle and the luminescence was recorded with the ChemiDoc™ XRS+ System from Bio-Rad and ImageLab version 4.0.1 software (Bio-Rad, Mississauga, ON, Canada).

### Arabidopsis susceptibility testing for salt and lincomycin

Arabidopsis seeds, collected from WT or UmPce3 lines grown concurrently, were sterilized and germinated on ½ MS medium, plain or supplemented with 150 mM NaCl or 40 µM lincomycin. The petri dishes were then placed horizontally in a growth chamber with 12 h light and 12 h dark conditions and the germination and growth of the seedlings were monitored. For salt treatment of older seedlings, 2-week-old soil grown seedlings were watered with 300 mM NaCl. The seedlings were watered with plain water to maintain the same alt concentration and their morphological phenotypes were monitored over the course of a few weeks.

### Enrichment and statistical analyses

Experiments in this study were performed in a minimum of three biological replicates. Average plus standard deviation is shown in all experiments, unless otherwise mentioned. The data were analyzed with *t*-tests, ANOVA with Tukey’s or other appropriate statistical procedures as indicated. Enrichment analyses were performed with the default settings of ShinyGO 0.85 and gprofiler (https://bioinformatics.sdstate.edu/go/; https://biit.cs.ut.ee/gprofiler/gost).

## Data availability

The mass spectrometry proteomics data from the isolated chloroplasts were deposited in the ProteomeXchange Consortium via the PRIDE partner repository with the dataset identifier PXD075117 (**Token:**eWyfMHlmZwXl) (Perez-Riverol et al., 2025). The data from the affinity purification-mass spectrometry experiments have also been deposited with the dataset identifier PXD074957.

## Author contributions

Conceptualization: D.D., M.K., J.W.K.; Formal analysis: D.D., M.K. K.T.D., K.S., O.V. J. G.- M.; Funding acquisition: V.L., J. G.-M., G.H.B., X.L., K.H., J.W.K.; Investigation: D.D., M.K. K.T.D., S.S., M.I., L.P., K.S., O.V. J. G.-M.; Project administration: J.W.K.; Supervision: M.K., V.L., O.V., G.H.B., X.L., K.H., J.W.K.; Validation: D.D., M.K. K.T.D., S.S., M.I., L.P., K.S., O.V. J. G.-M.; Visualization: D.D., M.K., K.S., J. G.-M.;Writing – original draft: D.D., M.K., K.S., J. G.-M., J.W.K. Writing – review & editing: D.D., M.K. K.T.D., S.S., M.I., L.P., V.L. K.S., O.V. J. G.-M., G.H.B., X. L., K.H., J.W.K.

## Acknowledgements Funding information

The authors thank Drs. Alice Barkan and Susan Belcher for generously providing seeds of maize mutants. This work was supported by a Discovery grant from the Natural Sciences and Engineering Research Council of Canada (to J.W.K.), by the NSERC-CREATE PRoTECT program, and by a four-year fellowship from the University of British Columbia (to D.Y.D.). J.W.K. is a Burroughs Wellcome Fund Scholar in Molecular Pathogenic Mycology, and the Power Corporation fellow of the CIFAR program: Fungal Kingdom Threats & Opportunities. X.L. holds a Tier I Canada Research Chair. We also acknowledge funding through the German Research Foundation (DFG) including an IRTG 2172 PRoTECT award (project ID 73134146), a DFG Heisenberg grant HE6977/3-1 (project ID 444679648) (to KH), Q ExactiveHF/U3000 funding (project ID INST 186/1230-1 FUGG (to Dr. Stefanie Pöggeler), and Core Facility funding (project ID INST 186/1465-1) (to OV).

## Supplemental Tables

**Supplemental Table S1. All identified proteins in the *Zea mays* chloroplast proteome during infection with *Ustilago maydis*.**

**Supplemental Table S2. Significantly different abundant proteins in the chloroplast proteome at the different time points or treatments.**

**Supplemental Table S3. GO term and keyword enrichment of the chloroplast proteome.**

**Supplemental Table S4. Candidate interacting proteins identified by affinity purification of UmPce3 and mass spectrometry with GO term analysis.**

**Supplemental Table S5. Primers, plasmids and strains.**

## Supplemental figures

**Supplemental Figure S1. Putative chloroplast localized effectors with chloroplast transit peptides from *U. maydis*.**

A) The average intensities from the proteomics data are shown for the proteins from infected samples minus the uninfected samples at 3, 5 and 7 dpi before and after proteinase K (PK) treatment. Standard deviation is shown and the data were analyzed with ANOVA plus Tukey’s test (p≤0.01 **). The red triangle depicts the timepoint and treatment where the protein was detected as differently abundant during the proteomics analysis (Supplemental Table S2).

B) The average gene transcript levels of the proteins in (A) during a time course of infection as reported by Lanver et al. 2018. The standard deviations are shown.

**Supplemental Figure S2. Alignment of the amino acid sequence of UMAG_05928 (UmPce3) with orthologs from other smut fungi, and with UMAG_05926 and UMAG_05927.**

A) Alignment with Clustal W of *U. maydis* UmPce3 with the orthologs identified in other smut fungi. The secretion signal peptide, as well as the location of the chloroplast transit pareptide are marked. Red squares and numbers denote conserved UmPce3 amino acid sequence regions between the different smut fungi including *Sporisorium reilianum* (SRS1_16551), *Pseudozyma hubeiensis* (PHSY_002000), *Ustilago trichophora* (UTRI_10400) and *Ustilago hordei* (UHO2_06808). B) Alignment with clustalW of UMAG_05926, UMAG_05927 andUMAG_05928 (UmPce3). The same amino acid stretches as in A are highlighted and numbered.

**Supplemental Figure S3. Confirmation of Umpce3 deletion in U. maydis strains 001 and 002 by genomic hybridization.**

A Southern blot was used to confirm the deletion of *Umpce3*. In the 001 strain background, isolates 61 and 62 are two independent mutants, while lane 67 represents the corresponding WT strain. In the 002 strain background, isolates 64 and 65 are two independent mutants while lane 68 represents the corresponding WT strain. For WT, a band of 957 bp is expected for the WT gene, and a band of 2524 bp is expected for the mutants after digestion of the genomic DNA with EcoRI and HindIII.

**Supplemental Figure S4. Impact of heterologous UmPce3 expression on *Arabidopsis* flowering, seed set and immunity.**

A) Days from planting until appearance of the first full flower for WT plants and the lines expressing UmPce3. The average plus standard deviation is shown. ANOVA plus Tukey’s test was used for statistical analysis (* p ≤ 0.05).

B) Percentage of sterile/aborted flowers which did not develop a seed pod for WT plants and lines expressing UmPce3.The average plus standard deviation is shown. ANOVA plus Tukey’s test was used for statistical analysis (*** p ≤ 0.001).

C) Number of seeds in a mature seed pod for WT plants and lines expressing UmPce3. The average plus standard deviation is shown. ANOVA plus Tukey’s test was used for statistical analysis (* p ≤ 0.05).

D) Number of conidia of *Hyaloperonospora arabidopsidis (Hpa*) Noco2 formed after 1 week of infection for WT, lines expressing UmPce3, and a mutant lacking EDS1 (compromised immunity). The average plus standard deviation is shown. ANOVA plus Tukey’s test was used for statistical analysis (* p ≤ 0.05; ** p ≤ 0.01).

E) Salicylic Acid measurement of *Arabidopsis* WT, UmPce3 expressing lines and the immunocompromised line EDS1. The different lines were not infected or infected with *Hpa* Noco2 and SA was measured. Average plus standard deviation is shown. ANOVA plus Tukey’s for statistic differences (* p ≤ 0.05; ** p ≤ 0.01; *** p ≤ 0.001; ns not significantly different).

**Supplemental Figure S5. Enlarged image of the volcano plot of UmPce3 interactors found in *Arabidopsis*.** A more detailed depiction of the volcano plot from Figure 4B shows the 382 identified putative UmPce3 interactors. A total of 279 significant positive interactors with p values of less than 0.05 were further analysed and marked in the quadrant “enriched in UmPce3” when the Log2 transformed LFQ protein intensity values of the UmPce3 plants compared to GFP expressing plants were p<0.05 according to a t-test, had a value of greater than 1 for t-test differences between those two lines and were detected in a second pulldown with HA tagged UmPce3. For more details, see Materials and Methods. Orange circles show proteins which were only detected in the UmPce3-GFP analysis while turquoise circles show proteins identified in the UmPce3-GFP as well as in the UmPce3-HA analysis. This identified 88 putative proteins including UmPce3. The circle sizes reflect the average MS/MS count for each depicted protein, and chloroplast proteins are marked with a black triangle. The effector UmPce3 (bait) and the putative interactor AtRH3 as best hits are labelled in the volcano plot and the threshold is indicated.

**Supplemental Figure S6. UmPce3-GFP peptides identified after affinity purification and mass spectrometry.**

A) Protein gel of the three independent UmPce3-2HA-mGFP5 expressing samples after affinity purification with anti-GFP beads. Colloidal Coomassie staining was employed.

B) Peptide location and intensity along the polypeptide sequence of UmPce3 for the peptides identified with IP-MS of UmPce3. Only peptides with low intensity were recorded upstream of the predicted cTP cleavage side (red dashed line).

**Supplemental Figure S7. Impaired chloroplast developmental interferes with biotrophy establishment and tumor formation in *Zea mays***

A) Disease symptoms observed in seedlings homozygous for the Zm *rh3b-1* allele and control plants. Seeds from segregating ears of rh3b-1/+ X rh3b-1/+ plants were planted and seedlings were infected with *U. maydis* at day 9. Disease symptoms were scored 2 weeks later and correlated with the plant genotypes determined by PCR (primers: Rh3b_for Rh3b_rev for WT (187 bp) and Rh3b_for and mu1/mu2 for transposon insertion in RH3b (157 bp)). Symptom class 5 shown in purple correlates to death of the 3^rd^ and/or 4^th^ leaf.

B) Disease symptoms observed for Zm *rh3b-1* and control plants. The observed distribution of disease symptoms for control plants and the Zm *rh3b-1* plants from A are shown together with the average observed disease index with standard deviation. The data were analyzed for statistical significance with a t-test with * p≤0.05.

C) Disease symptoms in Zm *ppr53* and control plants. Seeds from segregating ears of ppr53-2/+ X ppr53-3/+ were planted and seedlings were infected at day 9. Disease symptoms were scored at 2 weeks and correlated with the phenotype of defective chloroplast developmental in the Zm *ppr53* plants. Symptom class 5 shown in purple correlates to death of the 3^rd^ and/or 4^th^ leaves.

D) Disease symptoms in Zm *ppr53* and control plants. The observed distribution of disease symptoms for control and Zm *ppr53* plants from C are shown together with the average observed disease index with standard deviation. The data were analyzed for statistical significance with a t-test with *** p≤0.001. The seeds were generously provided by Dr. Alice Barkan.

**Supplemental Figure S8. Model for the impact of UmPce3 on maize during infection with *U. maydis*.**

The effector UmPce3 as a propeptide is secreted by *U. maydis* into the apoplast with cleavage of the signal peptide. The protein is further proteolytic processed after translocation from the apoplast into the plant cytosol and during translocation into the chloroplast. In the chloroplast, the mature active effector interacts with the chloroplast DEAD-box ATP dependent RNA helicase RH3 within the 30S/50S chloroplast ribosomal complex. This interaction is proposed to negatively impact chloroplast functions with implications for general plant growth and responses to abiotic and biotic stress (bold arrows). The effector further influences retrograde signaling from the chloroplast to the nucleus (bold arrow). Retrograde signaling induces gene expression changes in the nucleus in response to chloroplast disturbances. An influence on the expression of nuclear genes may contribute to the observed phenotypic changes (dashed arrows). During optimal plant growth, the effector UmPce3 is a virulence factor contributing to disease. In conditions of abiotic stress such as high salinity or activation of retrograde signaling by the antibiotic lincomycin, UmPce3 may dampen virulence. UmPce3 therefore may target chloroplast functions to support fungal biotrophy in a manner responsive to plant growth conditions.

