## Supplemental Figures S1-S8 for "A fungal effector targets the chloroplast to support biotrophy by balancing disease and plant health"

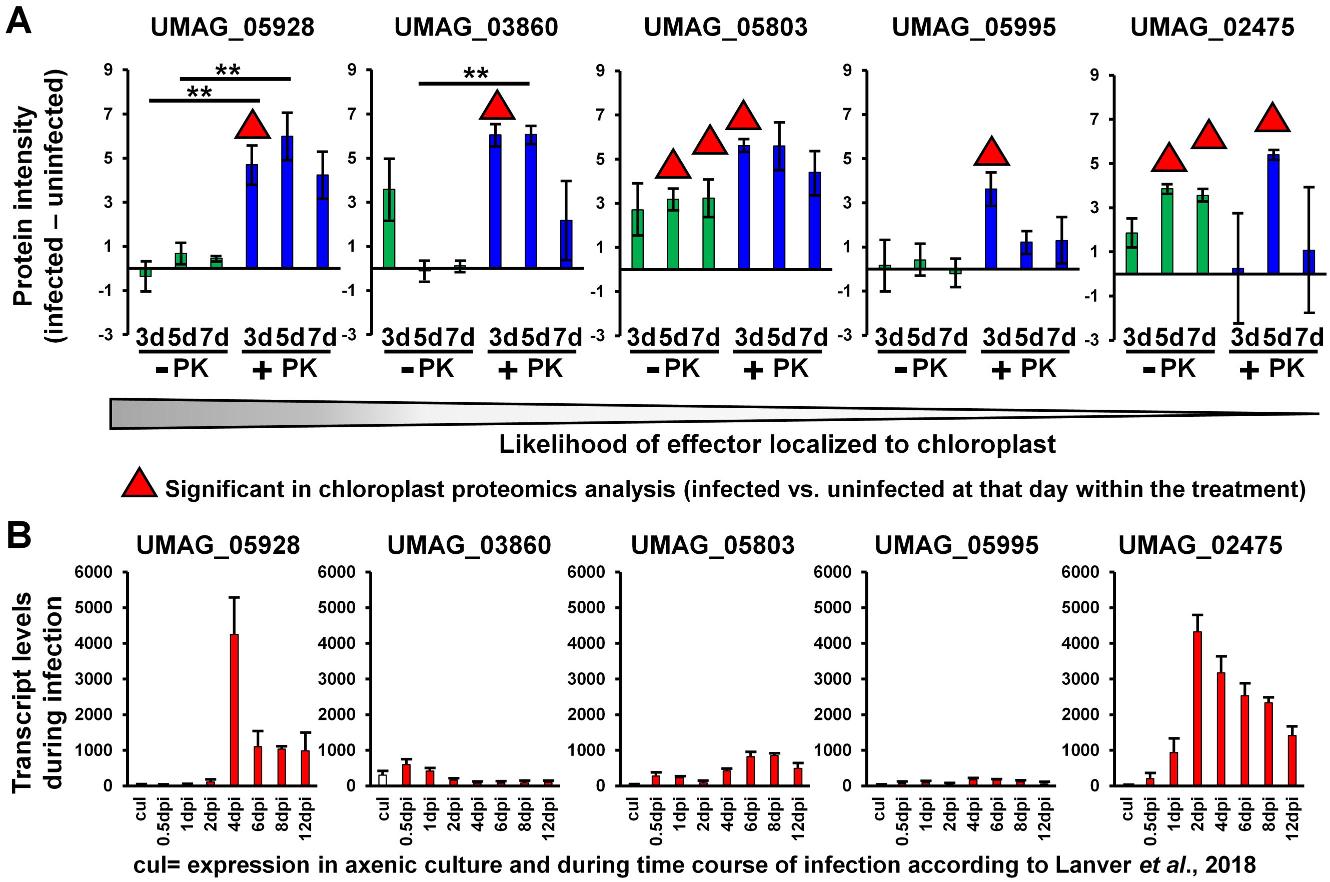


**Supplemental Figure S1. Putative chloroplast localized effectors with chloroplast transit peptides from *U. maydis*.**

A) The average intensities from the proteomics data are shown for the proteins from infected samples minus the uninfected samples at 3, 5 and 7 dpi before and after proteinase K (PK) treatment. Standard deviation is shown and the data were analyzed with ANOVA plus Tukey's test ($p \leq 0.01$ **). The red triangle depicts the timepoint and treatment where the protein was detected as differently abundant during the proteomics analysis (Supplemental Table S2).

B) The average gene transcript levels of the proteins in (A) during a time course of infection as reported by Lanver et al. 2018. The standard deviations are shown.

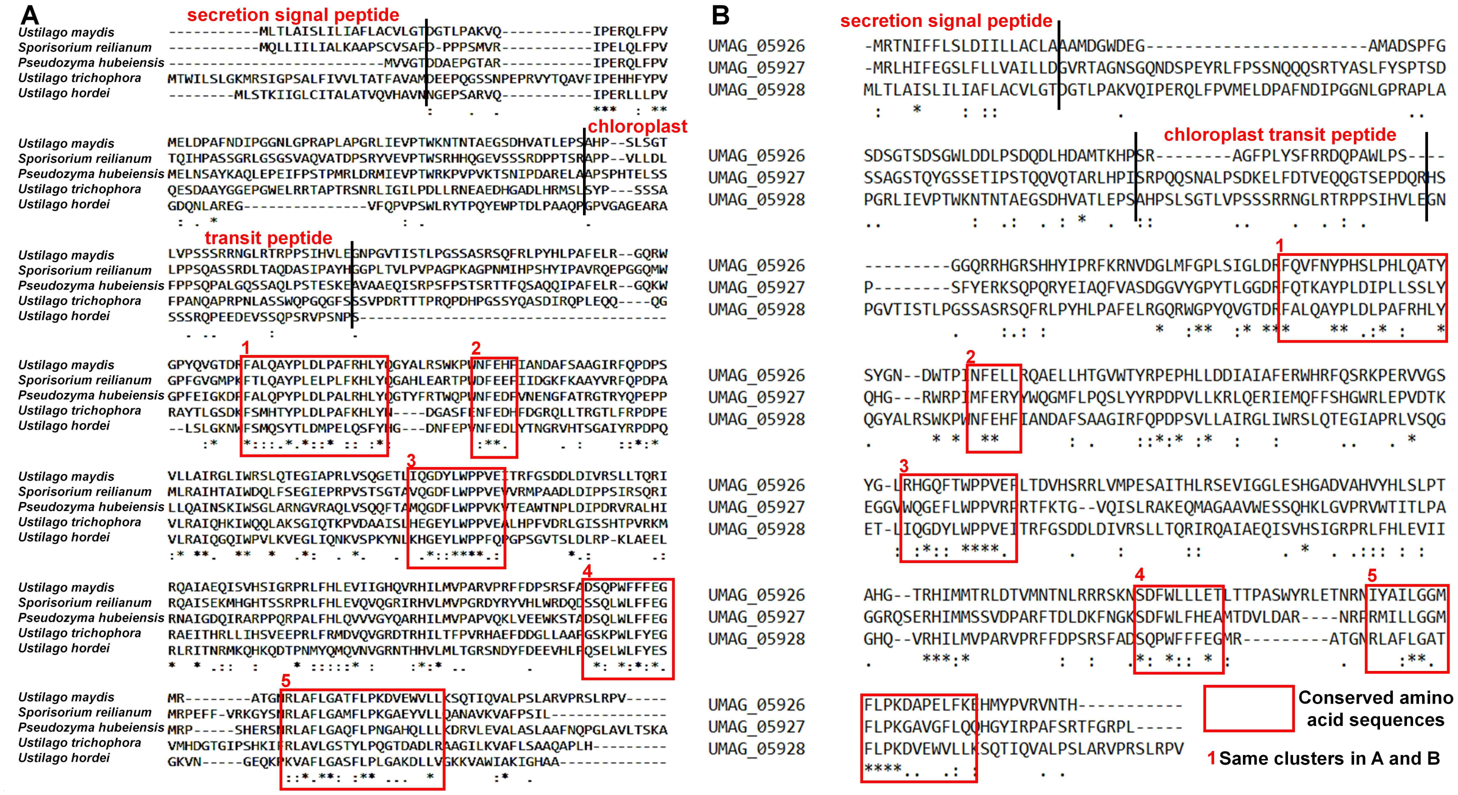


**Supplemental Figure S2. Alignment of the amino acid sequence of UMAG_05928 (UmPce3) with orthologs from other smut fungi, and with UMAG_05926 and UMAG_05927.**

A) Alignment with Clustal W of *U. maydis* UmPce3 with the orthologs identified in other smut fungi. The secretion signal peptide, as well as the location of the chloroplast transit pareptide are marked. Red squares and numbers denote conserved UmPce3 amino acid sequence regions between the different smut fungi including *Sporisorium reilianum* (SRS1_16551), *Pseudozyma hubeiensis* (PHSY_002000), *Ustilago trichophora* (UTRI_10400) and *Ustilago hordei* (UHO2_06808). B) Alignment with clustalW of UMAG_05926, UMAG_05927 andUMAG_05928 (UmPce3). The same amino acid stretches as in A are highlighted and numbered.

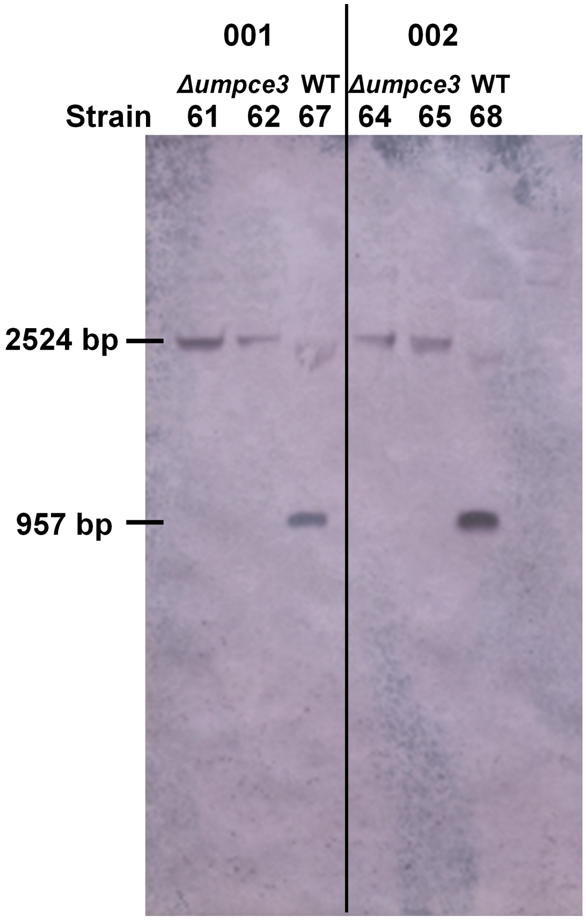


**Supplemental Figure S3. Confirmation of *Umpce3* deletion in *U. maydis* strains 001 and 002 by genomic hybridization.**

A Southern blot was used to confirm the deletion of *Umpce3*. In the 001 strain background, isolates 61 and 62 are two independent mutants, while lane 67 represents the corresponding WT strain. In the 002 strain background, isolates 64 and 65 are two independent mutants while lane 68 represents the corresponding WT strain. For WT, a band of 957 bp is expected for the WT gene, and a band of 2524 bp is expected for the mutants after digestion of the genomic DNA with EcoRI and HindIII.

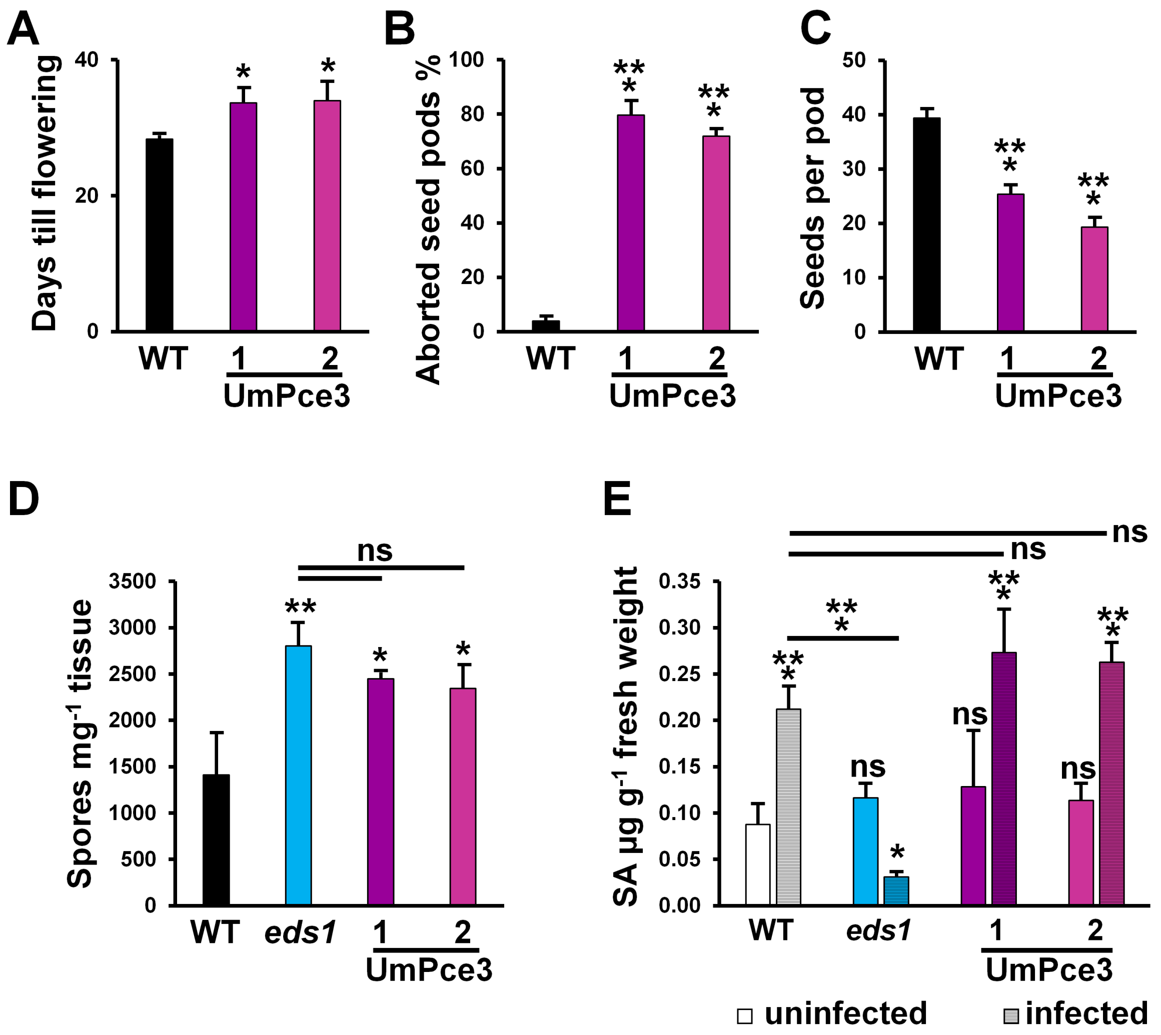


**Supplemental Figure S4. Impact of heterologous UmPce3 expression on *Arabidopsis* flowering, seed set and immunity.**

A) Days from planting until appearance of the first full flower for WT plants and the lines expressing UmPce3. The average plus standard deviation is shown. ANOVA plus Tukey's test was used for statistical analysis (* $p \leq 0.05$).

B) Percentage of sterile/aborted flowers which did not develop a seed pod for WT plants and lines expressing UmPce3.The average plus standard deviation is shown. ANOVA plus Tukey's test was used for statistical analysis (*** $p \leq 0.001$).

C) Number of seeds in a mature seed pod for WT plants and lines expressing UmPce3. The average plus standard deviation is shown. ANOVA plus Tukey's test was used for statistical analysis (* $p \leq 0.05$).

D) Number of conidia of *Hyaloperonospora arabidopsidis (Hpa*) Noco2 formed after 1 week of infection for WT, lines expressing UmPce3, and a mutant lacking EDS1 (compromised immunity). The average plus standard deviation is shown. ANOVA plus Tukey's test was used for statistical analysis (* $p \leq 0.05$; ** $p \leq 0.01$).

E) Salicylic Acid measurement of *Arabidopsis* WT, UmPce3 expressing lines and the immunocompromised line EDS1. The different lines were not infected or infected with *Hpa* Noco2 and SA was measured. Average plus standard deviation is shown. ANOVA plus Tukey's for statistic differences (* $p \leq 0.05$; ** $p \leq 0.01$; *** $p \leq 0.001$; ns not significantly different).

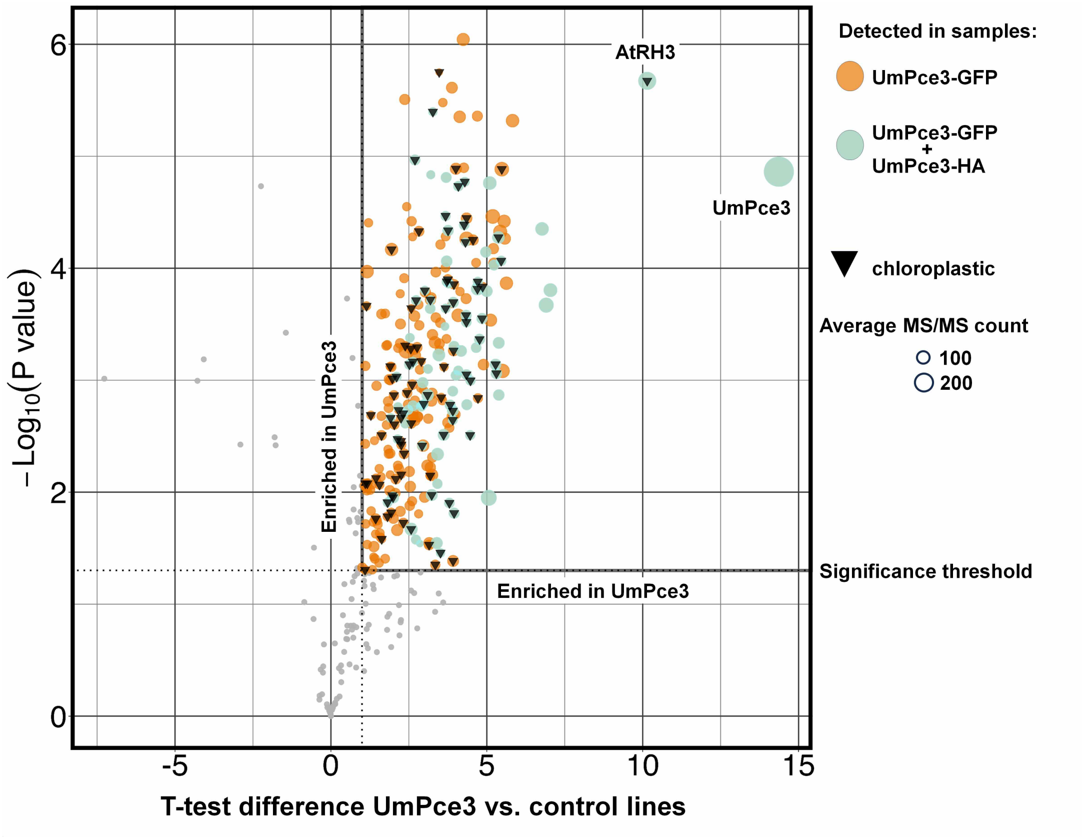


**Supplemental Figure S5. Enlarged image of the volcano plot of UmPce3 interactors found in *Arabidopsis*.** A more detailed depiction of the volcano plot from Figure 4B shows the 382 identified putative UmPce3 interactors. A total of 279 significant positive interactors with p values of less than 0.05 were further analysed and marked in the quadrant "enriched in UmPce3" when the Log2 transformed LFQ protein intensity values of the UmPce3 plants compared to GFP expressing plants were $p<0.05$ according to a t-test, had a value of greater than 1 for t-test differences between those two lines and were detected in a second pulldown with HA tagged UmPce3. For more details, see Materials and Methods. Orange circles show proteins which were only detected in the UmPce3-GFP analysis while turquoise circles show proteins identified in the UmPce3-GFP as well as in the UmPce3-HA analysis. This identified 88 putative proteins including UmPce3. The circle sizes reflect the average MS/MS count for each depicted protein, and chloroplast proteins are marked with a black triangle. The effector UmPce3 (bait) and the putative interactor AtRH3 as best hits are labelled in the volcano plot and the threshold is indicated.

A

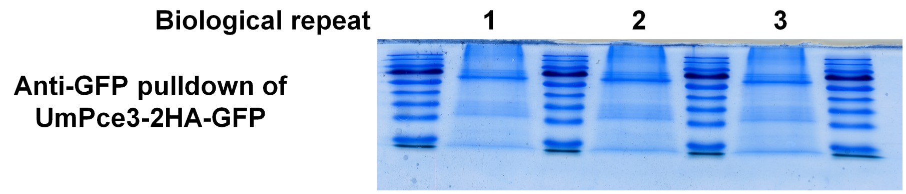


B

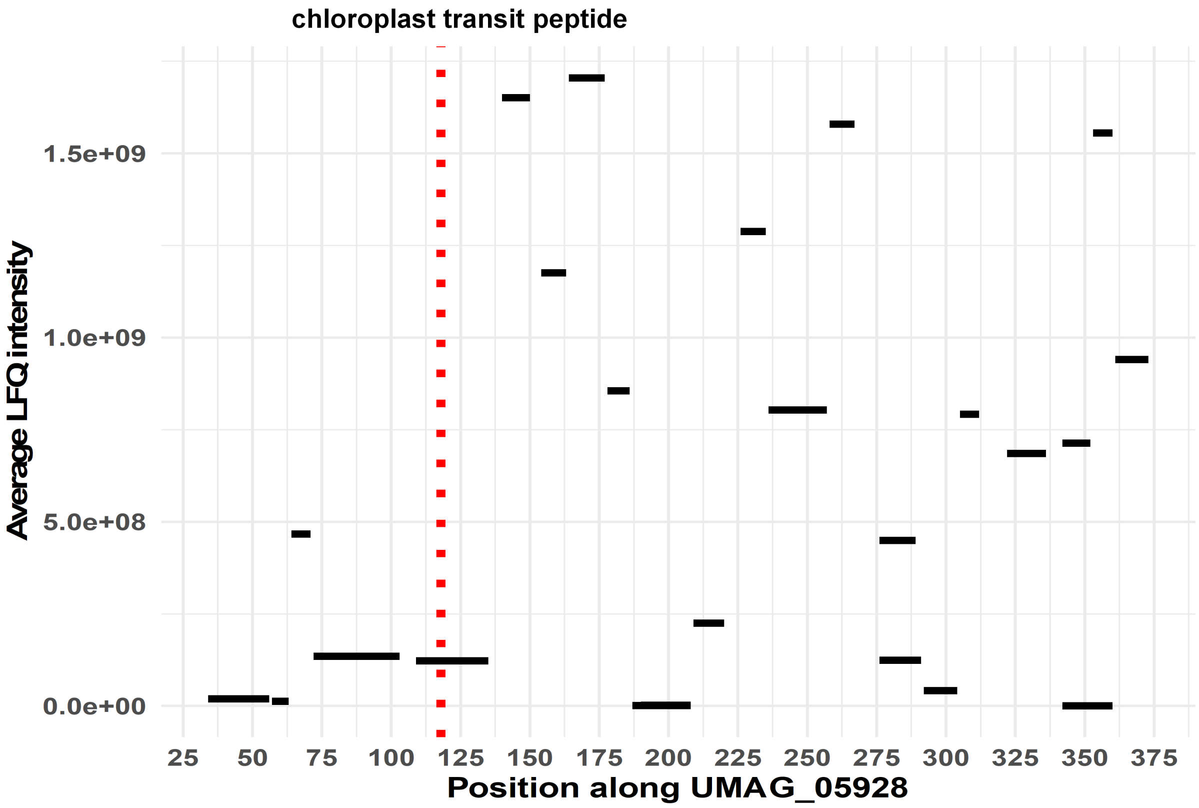


**Supplemental Figure S6. UmPce3-GFP peptides identified after affinity purification and mass spectrometry.**
A) Protein gel of the three independent UmPce3-2HA-mGFP5 expressing samples after affinity purification with anti-GFP beads. Colloidal Coomassie staining was employed.
B) Peptide location and intensity along the polypeptide sequence of UmPce3 for the peptides identified with IP-MS of UmPce3. Only peptides with low intensity were recorded upstream of the predicted cTP cleavage side (red dashed line).

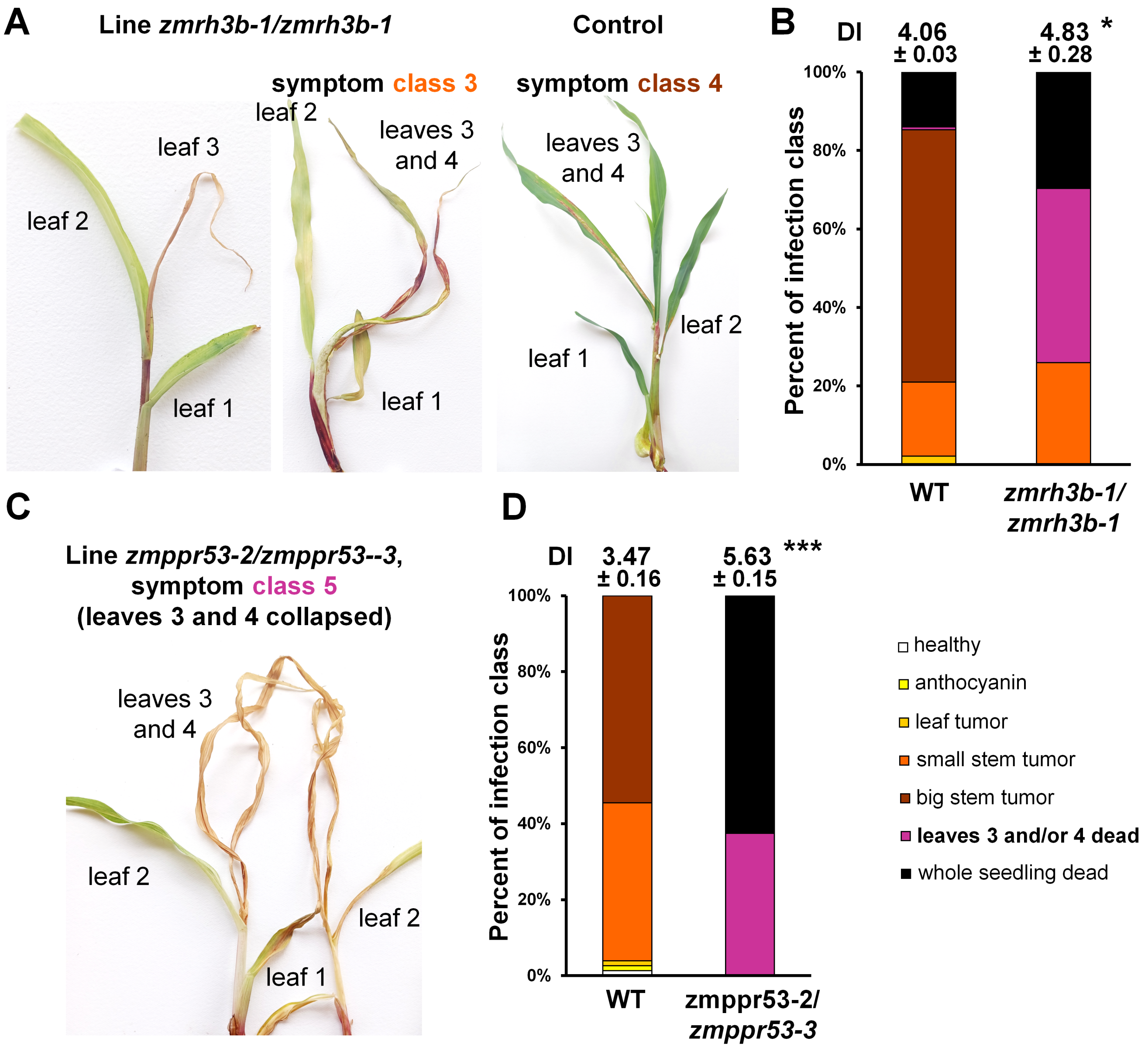


**Supplemental Figure S7. Impaired chloroplast developmental interferes with biotrophy establishment and tumor formation in *Zea mays***

A) Disease symptoms observed in seedlings homozygous for the Zm *rh3b-1* allele and control plants. Seeds from segregating ears of rh3b-1/+ X rh3b-1/+ plants were planted and seedlings were infected with *U. maydis* at day 9. Disease symptoms were scored 2 weeks later and correlated with the plant genotypes determined by PCR (primers: Rh3b_for Rh3b_rev for WT (187 bp) and Rh3b_for and mu1/mu2 for transposon insertion in RH3b (157 bp)). Symptom class 5 shown in purple correlates to death of the 3$^{rd}$ and/or 4$^{th}$ leaf.

B) Disease symptoms observed for Zm *rh3b-1* and control plants. The observed distribution of disease symptoms for control plants and the Zm *rh3b-1* plants from A are shown together with the average observed disease index with standard deviation. The data were analyzed for statistical significance with a t-test with * p≤0.05.

C) Disease symptoms in Zm *ppr53* and control plants. Seeds from segregating ears of ppr53-2/+ X ppr53-3/+ were planted and seedlings were infected at day 9. Disease symptoms were scored at 2 weeks and correlated with the phenotype of defective chloroplast developmental in the Zm *ppr53* plants. Symptom class 5 shown in purple correlates to death of the 3$^{rd}$ and/or 4$^{th}$ leaves.

D) Disease symptoms in Zm *ppr53* and control plants. The observed distribution of disease symptoms for control and Zm *ppr53* plants from C are shown together with the average observed disease index with standard deviation. The data were analyzed for statistical significance with a t-test with *** p≤0.001. The seeds were generously provided by Dr. Alice Barkan.

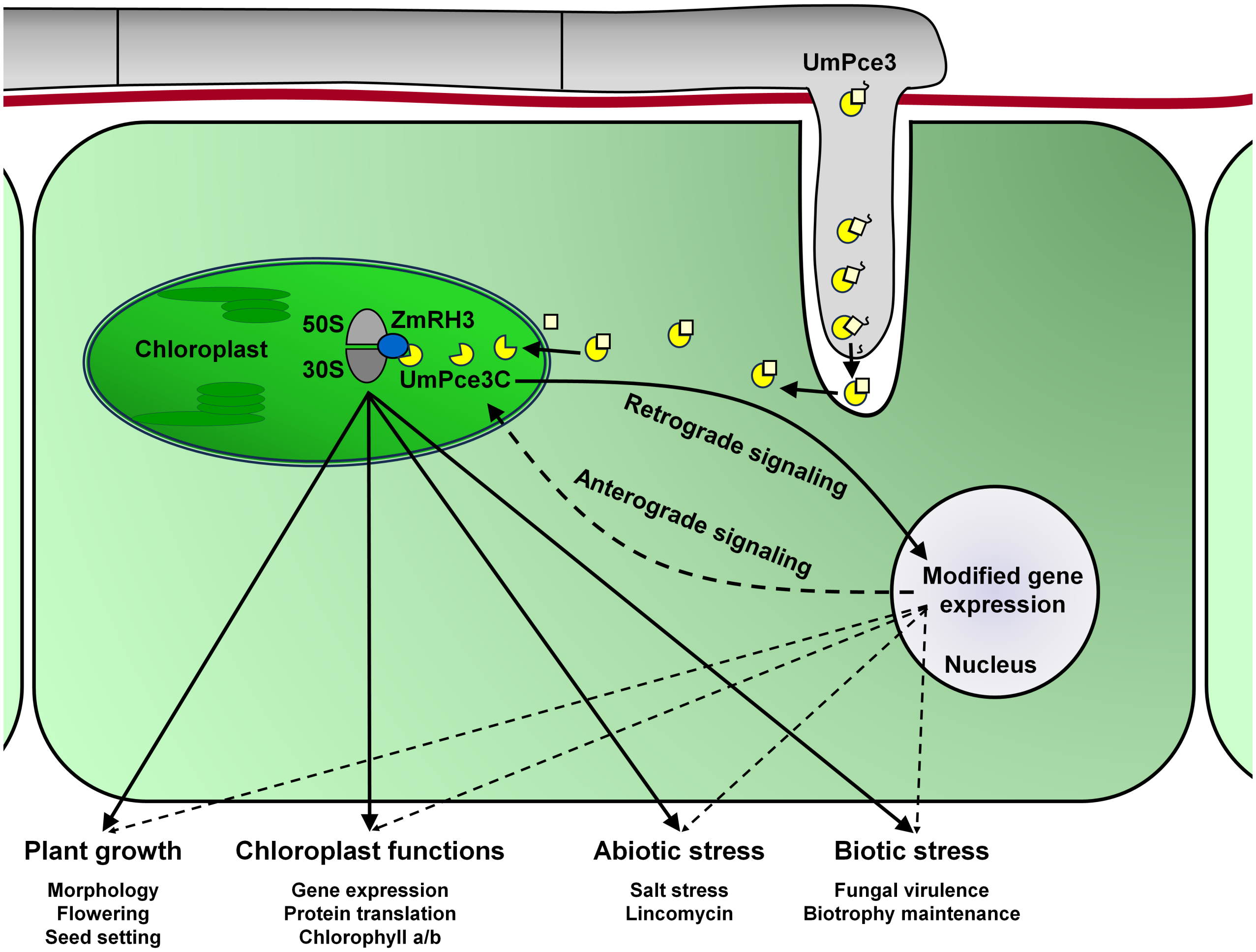


**Figure S8. Model for the impact of UmPce3 on maize during infection with *U. maydis*.**

The effector UmPce3 as a propeptide is secreted by *U. maydis* into the apoplast with cleavage of the signal peptide. The protein is further proteolytic processed after translocation from the apoplast into the plant cytosol and during translocation into the chloroplast. In the chloroplast, the mature active effector interacts with the chloroplast DEAD-box ATP dependent RNA helicase RH3 within the 30S/50S chloroplast ribosomal complex. This interaction is proposed to negatively impact chloroplast functions with implications for general plant growth and responses to abiotic and biotic stress (bold arrows). The effector further influences retrograde signaling from the chloroplast to the nucleus (bold arrow). Retrograde signaling induces gene expression changes in the nucleus in response to chloroplast disturbances. An influence on the expression of nuclear genes may contribute to the observed phenotypic changes (dashed arrows). During optimal plant growth, the effector UmPce3 is a virulence factor contributing to disease. In conditions of abiotic stress such as high salinity or activation of retrograde signaling by the antibiotic lincomycin, UmPce3 may dampen virulence. UmPce3 therefore may target chloroplast functions to support fungal biotrophy in a manner responsive to plant growth conditions.
